# Elevated MyoD1 levels expand genome-wide binding and the repertoire of regulated genes

**DOI:** 10.1101/2025.11.13.688391

**Authors:** Oscar N. Whitney, Gina M. Dailey, Joseph K. McKenna, Xavier Darzacq, Robert Tjian

## Abstract

Transcription factor (TF) upregulation accompanies many cellular state transitions, yet how increased TF abundance impacts gene regulation remains unclear. Two broad models are often invoked, whereby higher TF levels amplify the expression of pre-existing target genes, or, by mass-action binding, expand genome engagement and regulation to lower-affinity sites. We sought to elucidate how these two regulatory modes contribute to cell differentiation in a well characterized myogenic system by upregulating the expression of the myogenic TF MyoD1 in C2C12 myoblasts. Unexpectedly, elevated MyoD1 levels impaired myoblast fusion (a hallmark of myogenic differentiation), yet enabled robust contraction in myotubes that did form. Live-cell single-molecule imaging and CUT&RUN profiling revealed that elevated MyoD1 dosage increased total genome-wide chromatin binding and broadened genome occupancy by preferentially engaging lower-affinity sites. Integrating CUT&RUN with RNA-seq experiments linked expanded MyoD1 binding to upregulation of cell adhesion genes. Cell mixing and fractionated RNA-seq experiments supported a two-population model in which an adhesion-gene-upregulated, unfused myoblast population supported contraction of myotubes formed by fusion-competent cells. Ectopic expression of several individual MyoD1-upregulated cell adhesion genes was sufficient to recapitulate the “off script” myotube contraction phenotype. Together, these results support a MyoD1 dose-dependent “spillover” model, in which increased TF abundance broadens cis-regulatory engagement and produces distinct cell differentiation outcomes.

**Significance Statement:** Transcription factor dosage is a central control knob in many biological and disease-associated cell state transitions, but its range of gene regulatory consequences are poorly understood. By inducibly elevating MyoD1 expression in C2C12 cells, we uncovered a paradoxical outcome: elevated MyoD1 levels suppressed myoblast fusion yet enabled robust myotube contraction. Elevated MyoD1 levels expanded MyoD1 genome occupancy through engagement of low-affinity sites and upregulated adhesion genes whose individual ectopic expression was sufficient to recapitulate the contraction phenotype. Mixing experiments indicated a non-cell-autonomous contribution from an adhesion-gene-high subpopulation that supported contraction of myotubes. Our results demonstrate that TF dosage can expand regulatory scope and yield qualitatively different differentiation outcomes.

## Introduction

Developmental cell state transitions are driven by the expression and upregulation of cell lineage-specific transcription factors (TFs).^1–4^ TF upregulation frequently accompanies cell fate decisions, and ectopic expression of some TFs can be sufficient to reprogram cells towards TF-dependent identities.^5–8^ Despite this central role, the mechanisms by which increases in TF dosage alter genome engagement and transcriptional output to specify differentiation outcomes remain incompletely defined.

Classical studies employing *in vitro* transcription reconstitution and *in cellulo* transcription reporter assays have reported that increased TF dosage can amplify the expression of target genes bearing high-affinity DNA motifs.^9–12^ These findings have led to the intuitive view that increased cell-type specific TF dosage largely serve to amplify target gene expression to reinforce TF-associated cell identity.^2,13^ However, mass-action binding predicts that as TF dosage increases, occupancy at high-affinity sites would approach saturation and progressively extend to lower-affinity “non-cognate” loci, potentially broadening a TF’s regulatory repertoire.^14,15^ Whether TF upregulation in differentiating cells primarily amplifies a defined target program, or instead expands binding to additional targets, and how these modes might be coupled to phenotypic outcomes remains unclear.

To interrogate TF dose-dependent regulatory outcomes in a tractable model of TF-dependent differentiation, we inducibly elevated expression of the TF MyoD1 during myogenic differentiation of murine C2C12 myoblasts.^16^ MyoD1 is necessary for muscle differentiation, which culminates in myoblasts’ fusion into multi-nucleated myotubes.^3^ Ectopic MyoD1 expression is also sufficient to reprogram diverse somatic cell types towards a skeletal muscle cell identity.^5^ In both contexts, MyoD1 has been reported to bind high-affinity E-box motifs in the cis-regulatory elements of genes required for myogenesis, such as *Myogenin*, thereby activating a cascade of genes that drive a myogenic differentiation program.^17,18^ However, whether increased MyoD1 dosage re-distributes genome occupancy between high-affinity myogenic targets and lower-affinity sites, and how such redistribution might impact transcription regulation and differentiation outcomes remains poorly explored.

In this report, we tested how synthetic elevation of MyoD1 shapes genome engagement and transcriptional regulation during C2C12 differentiation. Using phenotypic assays, single molecule tracking, and CUT&RUN profiling combined with RNA sequencing, we show that elevated MyoD1 dosage surprisingly impaired myoblast fusion but enabled myotube contraction. This paradoxical phenotype was underpinned by a non-cell-autonomous mechanism consistent with two cell populations, in which unfused myoblasts activated an “off-script” adhesion gene program that supported contraction of myotubes formed by fusion-competent cells. Mechanistically, increased MyoD1 abundance broadened genome occupancy via the engagement of lower-affinity targets, expanding the MyoD1 regulatory repertoire to promote an alternative transcriptional program.

## Results

### Elevated MyoD1 levels inhibit myoblast fusion but enable myotube contraction

To manipulate TF dosage during a cell state transition, we engineered murine C2C12 myoblasts to express a doxycycline inducible 3xFlag-Halo-MyoD1 transgene. Subsequently, we tagged the endogenous MyoD1 locus with N-terminal 3xFlag-Halo (C2C12^MyoD1::Halo + iMyoD1^), enabling us to observe the expression dynamics of the entire MyoD1 protein pool **(Fig. S1 A-E)**. Dox induction of Halo-MyoD1 at the onset of C2C12^MyoD1::Halo + iMyoD1^ differentiation **(Fig. 1A, B)** increased total nuclear Halo-MyoD1 expression in expressing nuclei by 2.88-fold after 6 h of dox treatment and differentiation, 1.77-fold after 24 h and returned to near endogenous levels by 48 h **(Fig. 1C,D)**. Dox treatment also increased the fraction of nuclei expressing Halo-MyoD1 by ∼40% at 6 and 24 h of differentiation **(Fig. S1F)**. To minimize potential clone-specific effects from genome editing and clonal isolation, we also generated a polyclonal C2C12 population bearing additional dox-inducible V5-Halo-MyoD1 alleles (C2C12^iMyoD1^). C2C12^iMyoD1^ exhibited a comparable induction fraction to the clonal C2C12^MyoD1::Halo + iMyoD1^ line (∼30% Halo-MyoD1-positive nuclei at 24 h) **(Fig. S2B)**. We next asked whether induced MyoD1 dosage could be tuned by dox titration. Dox concentration primarily modulated the fraction of C2C12^iMyoD1^ that expressed Halo-MyoD1, whereas MyoD1 abundance within induced nuclei was similar **(Fig. S2E-F)**, which has been previously observed in other dox-inducible systems.^19,20^ Accordingly, subsequent analyses compare untreated (endogenous MyoD1 levels) and dox-treated (elevated MyoD1 levels) conditions, rather than a continuous per-cell dosage series.

**Figure 1.**
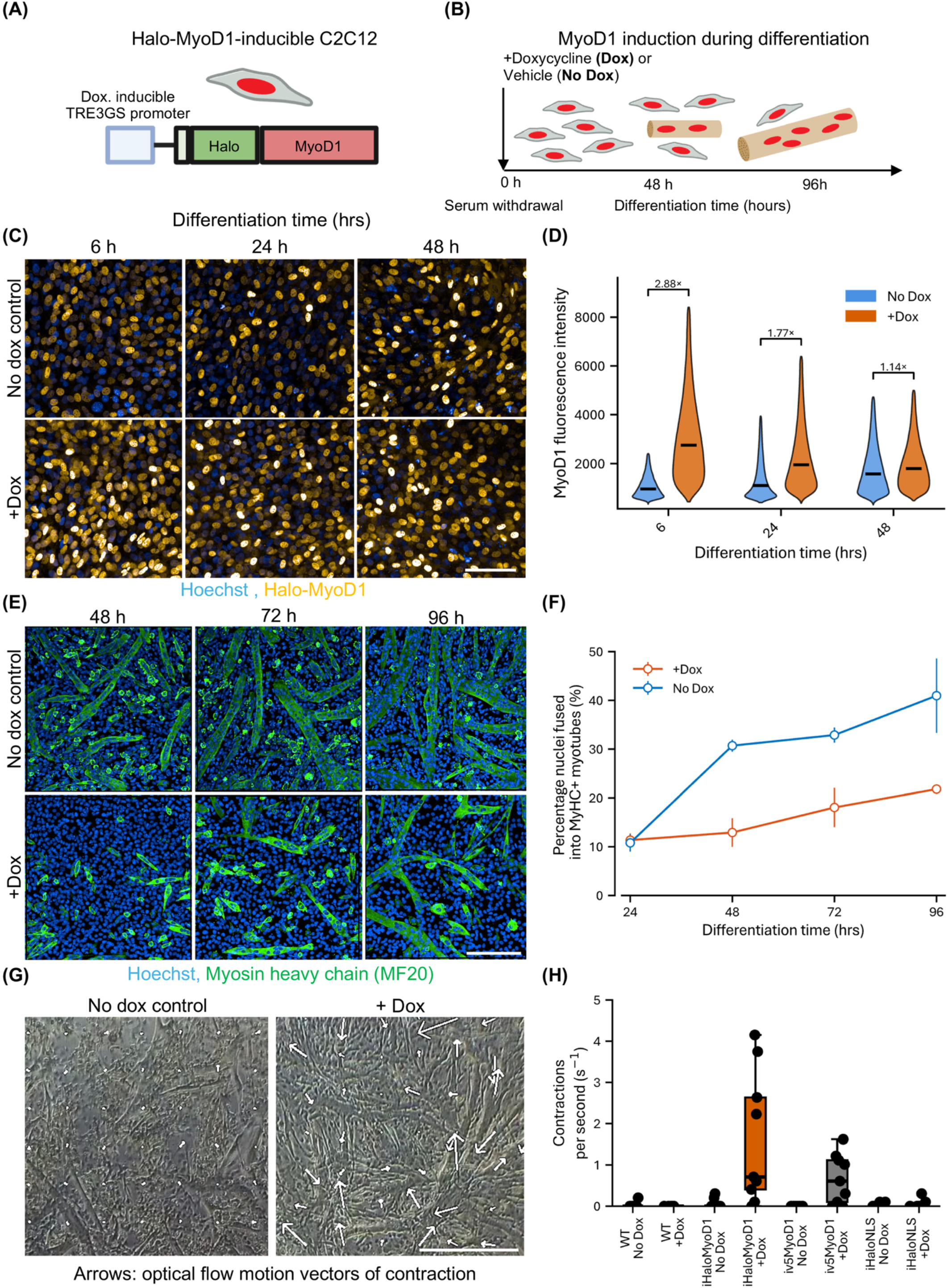
Elevated MyoD1 levels inhibit myoblast fusion but enable myotube contraction. (A) Schematic describing Halo-MyoD1-inducible C2C12: C2C12 myoblasts (cell with red nucleus) containing doxycycline-inducible TRE3GS promoter-driven (white box) V5 or 3xFLAG-Halo-tagged *MyoD1* alleles. (B) Schematic describing Halo-*MyoD1* induction during myogenic differentiation: MyoD1 was induced via +dox treatment simultaneous with the onset of myogenic differentiation, which was triggered via serum withdrawal. Schematic also indicates C2C12 myoblast (grey cells/red nucleus) fusion into myotubes (multinucleated tube) at 48 h, and maximal myotube fusion at 96 h. (C) Representative images of C2C12^MyoD1::Halo + iMyoD1^ cell line (A) treated with +dox or no dox over 48 h of differentiation. Yellow: JFX549 Halo-tag ligand-labeled Halo-MyoD1. Blue: Hoechst labeled nuclei. Scale bar: 100µm. (D) Violin plot quantification of C2C12^MyoD1::Halo + iMyoD1^ single-nucleus Halo-MyoD1 mean fluorescence intensity at 6, 24 and 48 h of differentiation ± dox treatment. Violins represent distribution of per-nucleus intensities pooled over 3 biological replicates; black lines indicate medians per condition/timepoint. Blue violins represent Halo-MyoD1 intensity of untreated (no dox) myoblasts, orange violins represent dox treated myoblasts. Analysis represents a minimum of 100,000 nuclei analyzed per time point and condition over three biological replicates. (E) Representative confocal microscopy images of C2C12^iMyoD1^ culture nuclei (Hoechst: blue) and myotubes (Myosin Heavy Chain immunofluorescence: green) over 48 - 96 h differentiation ± dox. Scale bar: 200 µm. (F) Quantification of ± dox treated C2C12^iMyoD1^ nuclei fusion into MyHC-expressing myotubes (as represented in Figure 1E). Orange line represents the fusion of +dox treated C2C12^iMyoD1^ , while blue line represents untreated C2C12^iMyoD1^. Plotted dots and bars represent means and standard deviation across 3 biological replicates. Analysis represents a minimum of 100,000 nuclei analyzed per time point and condition over three biological replicates. (G) Phase-contrast microscopy snapshots of C2C12^iMyoD1^ timelapse after 144 h of differentiation ± dox treatment. White arrows denote optical flow vectors of frame-to-frame pixel movement as a measurement of myotube contraction. Scale bar: 200 µm. (H) Boxplot quantification of myotube contraction: Contractions per second in C2C12^iHaloMyoD1^, C2C12^WT^, C2C12^iV5MyoD1^ and C2C12^iHaloNLS^ treated ± dox and differentiated for 144 h. Dots are well replicates performed across 3 biological replicates; line represents median of well replicates, box edges represent quartile 1 (lower) and quartile 3 (upper) , whiskers represent 1.5× IQR.

To assess the impact of elevated MyoD1 levels on myogenic differentiation, we treated C2C12^iMyoD1^ with dox at the onset of serum withdrawal-induced differentiation. Dox-treated C2C12^iMyoD1^ exhibited a ∼2-fold reduction in fusion at 48 - 96 h of differentiation relative to untreated controls **(Fig. 1E, F)**. This impairment in fusion was recapitulated in C2C12 bearing dox-inducible V5-MyoD1 alleles, controlling for HaloTag related effects **(Fig. S3C)**. Dox treatment of wild-type C2C12 (C2C12^WT^), and induction of Halo-NLS protein (C2C12^iHaloNLS^) did not affect fusion, controlling for doxycycline and protein induction-related effects, respectively **(Fig. S3A-C)**. Consistent with reduced fusion, Halo-MyoD1-expressing myoblasts were also approximately half as likely to simultaneously express MyoG, a marker of myogenic differentiation, at 24 h of differentiation **(Fig. S2A**,**C)**.^21^ Within single cells, increasing Halo-MyoD1 dosage anticorrelated with the likelihood of Myogenin expression, both at 24 h and 48 h of differentiation, indicating that elevated MyoD1 expression impaired commitment to myogenic differentiation **(Fig. S2D)**.

Despite impaired fusion, the myotubes that formed within dox-treated C2C12^iMyoD1^ cultures exhibited spontaneous and sustained contractions after 144 h of differentiation **(Fig. 1G, H, Supplementary Video 1)**. Under the same culture and imaging conditions, contractions were not observed in untreated C2C12^iMyoD1^, dox-treated C2C12^WT^, or dox-treated C2C12^iHaloNLS^ controls **(Fig. 1G,H)**. Contractions were also observed in C2C12 bearing dox-inducible V5-MyoD1 alleles (C2C12^iV5MyoD1^), controlling for HaloTag fusion-related effects **(Fig. 1H)**. While spontaneous C2C12 myotube contraction has been previously reported, it is uncommon, dependent on electrical pulse stimulation or culture on adhesion-promoting substrates.^22,23^ We never observed spontaneous contraction of C2C12^WT^ under our experimental conditions. As elevated MyoD1 levels impaired canonical differentiation outputs (fusion and *MyoG* activation) yet produced robust contraction of the myotubes that were able to form, we next asked whether increased MyoD1 dosage amplified the canonical myogenic transcriptional program or instead activated an alternative, dose-dependent gene expression program underlying this phenotype.

### Elevated MyoD1 levels do not amplify myogenic genes but upregulate an alternate transcriptional program

We initially sought to test whether increased MyoD1 dosage amplified high affinity target genes normally upregulated during myogenesis or instead enabled regulatory expansion via mass-action binding to an alternate set of target genes that would ordinarily not be expressed or be expressed at low levels. To test this hypothesis, we assessed the impact of elevated MyoD1 expression on the transcriptomes of undifferentiated (treated ±dox for 24 h in serum-rich growth media) and differentiating myoblasts (24 h differentiation and simultaneous ±24 h dox) in C2C12^iMyoD1^ **(Fig. 2A)**. MyoD1 induction affected gene transcription in both proliferating (336 up, 198 down) and 24 h differentiating myoblasts (992 up, 784 down) **(Fig. 2B,C)**.

**Figure 2.**
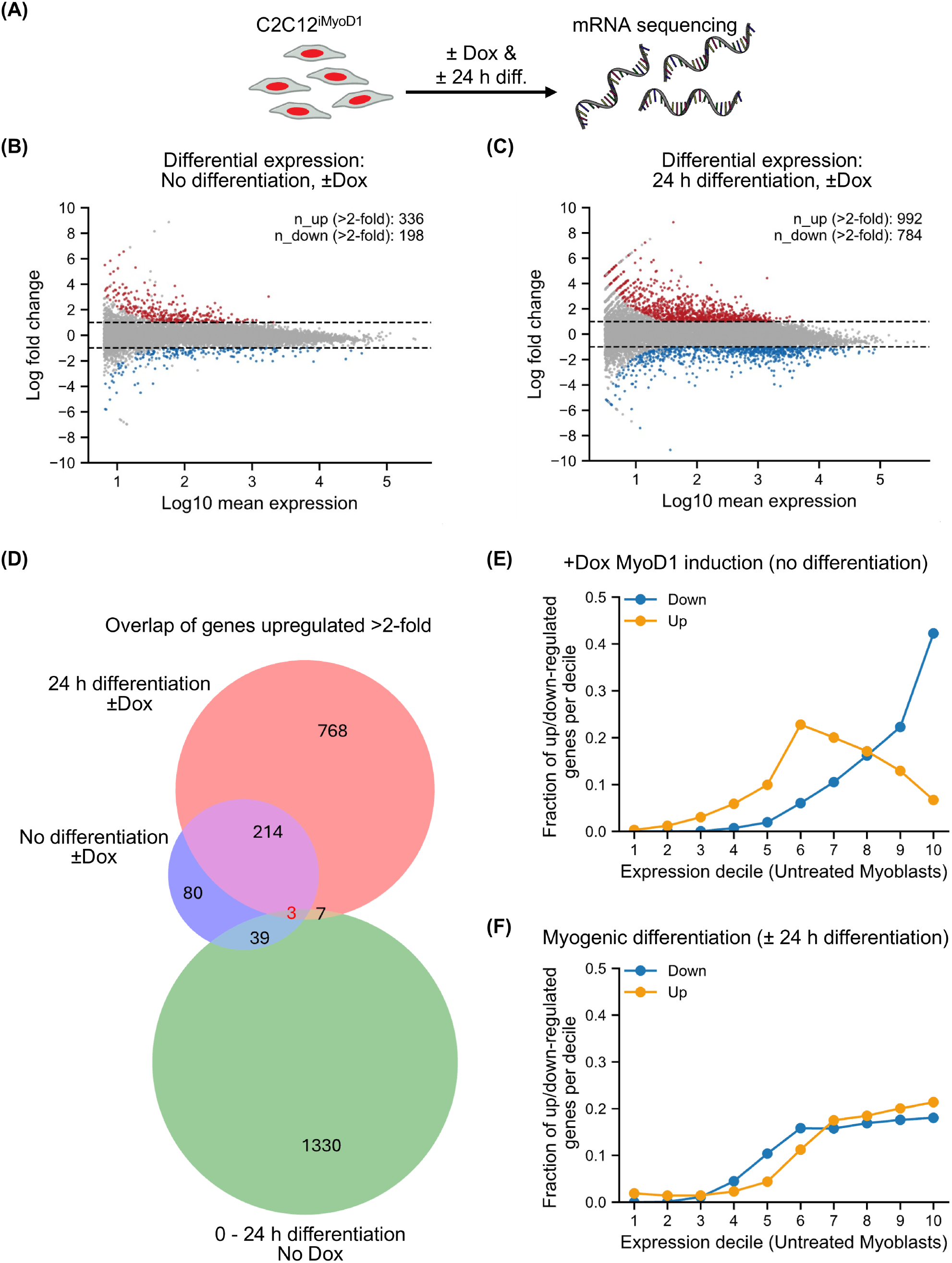
Elevated MyoD1 levels do not amplify myogenic genes but upregulate an alternate transcriptional program. (A) Schematic describing mRNA sequencing of C2C12^iMyoD1^ following ± dox treatment and ± 24 h differentiation. Grey/red cells represent C2C12^iMyoD1^, single-stranded nucleic acid represents RNA. (B) Gene expression log-ratio to average (MA) plot of C2C12^iMyoD1^ RNA-seq. ± dox and no differentiation. Red dots indicate significantly (p. adj<0.05) >2-fold upregulated genes. Blue dots indicate significantly (p. adj<0.05) >2-fold downregulated genes. Dashed lines represent a 2-fold change threshold. N_up and N_down represent the number of genes up/down-regulated as described above. (C) MA plot of C2C12^iMyoD1^ RNA-seq. ± dox and 24 h differentiation, as in (B). (D) 3-way Venn diagram of upregulated gene overlap (>2-fold, p.adj<0.05) between genes upregulated due to +dox treatment without differentiation (Blue circle), +dox treatment with 24 h differentiation (Red circle) and due to early endogenous myoblast differentiation (0-24 h differentiation, no dox, green circle). Number annotation represents the number of upregulated genes in each Venn category. (E) Distribution of significantly up-or downregulated genes due to +dox induction as compared to those genes’ basal TPM expression decile in untreated, proliferating myoblast RNA seq. (F) Distribution of significantly up-or downregulated genes due to 0-24 h of myogenic differentiation, as in (E).

To test whether MyoD1 induction amplified the endogenous myogenic transcriptional program, we compared dox-induced genes (MyoD1 dose-responsive genes) to genes upregulated due to early and later differentiation intervals (0–24, 24–48, and 48–120 h differentiation). Only ∼4% of dox-upregulated genes overlapped with genes upregulated within 24 h of native differentiation **(Fig. 2D)**, with modest overlap at later differentiation intervals (∼16% for 24–48 h; ∼9% for 48–120 h; **Fig. S4B,C)**. Expression of a curated panel of genes upregulated during early native myogenic differentiation, many of which are reported Myod1 targets, showed that dox treatment of proliferating C2C12^iMyoD1^ cells did not precociously activate the expression of myogenic genes, such as Myogenin, sarcomere components, or the fusogens Myomaker/Myomixer **(Fig. S4A)**.^5,17,24,25–30^ Instead, we observed that at 48 h of differentiation, dox treatment downregulated many myogenic genes in C2C12^iMyoD1^, including *MyoG* and the fusogenic peptides *Mymk/Mymx* **(Fig. S4A)**. Transcriptome-wide analysis supported this result, showing that all genes upregulated due to 24-48 h of endogenous differentiation were slightly but significantly downregulated by dox treatment, as compared with all other genes **(Fig. S4D)**. Genes upregulated by endogenous differentiation (0-24, 24-48 and 48-120 h of differentiation) enriched gene ontology terms such as myofibril, contractile muscle fiber and sarcomere **(Fig. S4E)**. In contrast, dox-induced genes primarily enriched cell membrane-related processes, including GO terms such as cell projection and cell junction, but did not enrich myofibril, contractile muscle fiber or sarcomere terms **(Fig. S4E)**. These results further suggest that the MyoD1-dose responsive program is distinct, rather than an amplification of the endogenous myogenic differentiation program.

Given these findings, we suspected that increased MyoD1 dosage may preferentially upregulate genes that were otherwise lowly expressed during myoblast differentiation. To test this possibility, we assessed MyoD1-dose responsive genes’ starting expression level in untreated, proliferating myoblasts. This analysis revealed that increased MyoD1 dosage preferentially upregulated genes in intermediate expression deciles (decile 6) while downregulating genes in higher expression deciles (decile 10) **(Fig. 2E)**. Serum-starvation-mediated myogenic differentiation preferentially up- and downregulated genes within the highest expression deciles (7-10) **(Fig. 2F)**. This result suggests that increased MyoD1 dosage upregulated normally lowly expressed genes, rather than amplifying the expression of highly expressed genes. We next sought to illuminate the gene regulatory mechanisms underlying dose-dependent MyoD1 activation of an alternative regulatory program.

### Elevated MyoD1 levels expand genome-wide binding via lower-affinity site engagement

In accordance with mass-action mediated genomic binding, we hypothesized that increased MyoD1 dosage may enable binding of additional chromatin sites to activate an alternate gene regulatory program. To assess MyoD1 chromatin binding in live cells, we employed single-molecule microscopy and tracking (SMT). Previous studies have linked slow single-molecule TF diffusion (<0.1 µm^2^/s) to chromatin binding.^31,32^ Therefore, we tested the impact of MyoD1 dosage on the fraction of Halo-MyoD1 molecules diffusing slower than 0.1 µm^2^/s (chromatin bound). We measured the diffusion dynamics of Halo-MyoD1 in our endogenously tagged and inducible Halo-MyoD1 (C2C12^MyoD1::Halo + iMyoD1^) line treated with dox or differentiated via serum starvation to increase total MyoD1 levels **(Fig. 3A)**. We measured and correlated paired MyoD1 fluorescence intensity and chromatin bound fractions in single myoblasts and observed that the MyoD1 chromatin bound fraction did not appreciably change over increasing MyoD1 mean fluorescence intensity, remaining near a 50% bound fraction over a 4.5-fold dosage increase **(Fig. 3B)**. A constant bound fraction with increased MyoD1 abundance suggests that the absolute number of chromatin-bound MyoD1 molecules increases with dosage. Thus, increased MyoD1 abundance scales genome engagement in a manner consistent with a mass-action regime.

**Figure 3.**
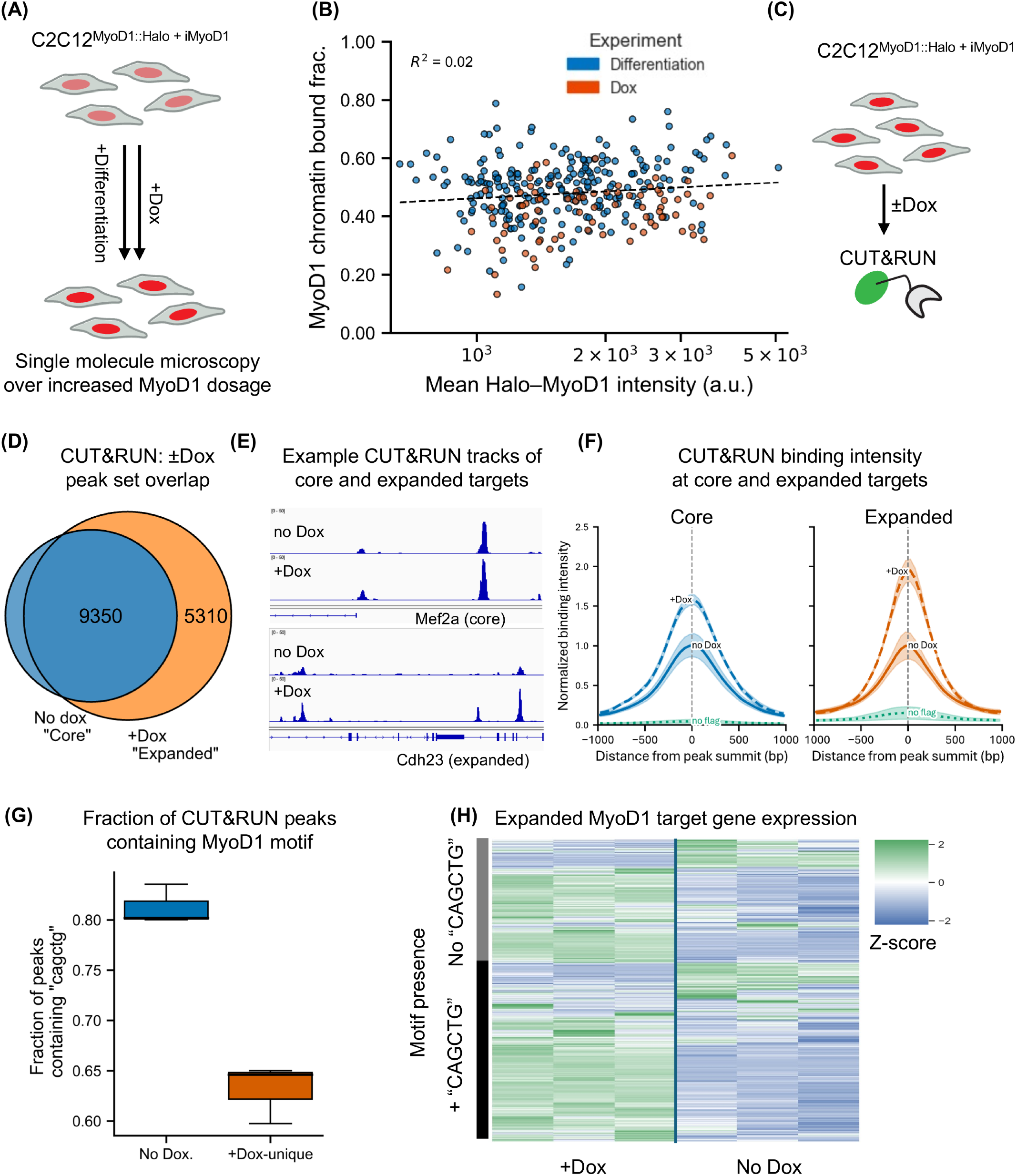
Elevated MyoD1 levels expand genome-wide binding via lower-affinity site engagement. (A) Schematic describing increased Halo-MyoD1 dosage in C2C12^MyoD1::Halo +^ iMyoD1 (cells with red/pink nuclei) due to + dox treatment or differentiation. (B) Scatterplot of all paired single-nucleus Halo-MyoD1 intensity and chromatin bound fraction measurements (fraction of Halo-MyoD1 molecules diffusing <0.1 µm^2^/s). Scatterplot includes all nuclei over ± dox treatment (orange dots) and differentiation time points (0, 24, 48, 72 h, blue dots). Dashed line represents best-fit linear regression on fluorescence intensity vs. bound fraction. R^2^ represents the proportion of MyoD1 bound fraction variance explained by MyoD1 fluorescence intensity. (C) Schematic describing CUT&RUN and sequencing of C2C12^MyoD1::Halo + iMyoD1^ cultures (cells with red nuclei) ± dox induction. Cells with red nuclei describe C2C12^MyoD1::Halo + iMyoD1^. Green oval and arm represent anti-flag antibody and MNase enzyme employed for CUT&RUN. (D) Venn diagram overlap of C2C12^MyoD1::Halo + iMyoD1^ ±dox CUT&RUN peak sets. Peak set represents peaks reproducible in at least 2/3 biological replicates. Blue circle defines “Core” targets bound at endogenous MyoD1 dosage. Orange circle defines “Expanded” targets bound at dox-induced dosage. Numerical annotations define number of Core and unique Expanded binding peaks. (E) Example IGV genome viewer tracks of Halo-MyoD1 CUT&RUN coverage ± dox at “core” gene Mef2a and “expanded” gene Cdh23. (F) Distribution of CUT&RUN signal coverage over no dox (Core targets) peak set and +dox (Expanded targets). No dox signal intensity was set to 1, to represent relative +dox signal intensity change. Dashed lines indicate +dox signal intensity, solid lines represent no dox signal intensity, and dotted green line indicates no flag control (anti-flag CUT&RUN performed in C2C12^WT^). Shading indicates standard deviation of signal over three biological replicates. (G) Fraction of no-dox and +dox-unique peak set containing MyoD1 consensus sequence “CAGCTG”. Motif presence was quantified in 100bp windows extended on each side of peaks in each peak set. Boxplots represent 3 biological replicates, where line indicates median, whiskers: 1.5x inter-quartile range, box edges: 1^st^ and 3^rd^ quartiles. (H) Heatmap of RNA expression (TPM, row z-scored) for genes linked to +Dox-unique MyoD1 CUT&RUN peaks. Genes were split by motif presence (black, +”CAGCTG”) or absence (grey, no “CAGCTG”) of a MyoD1 consensus CAGCTG sequence within their associated peaks. Rows represent peak-associated gene expression, clustered within each motif. Columns represent three biological replicates each of untreated and +dox treated C2C12^iMyoD1^ RNA-seq (as described in Fig. 2 A). Color scale indicates relative expression (blue: downregulated, green: upregulated).

We asked whether this increase in total MyoD1 binding occurs via enhanced binding of high- or lower-affinity chromatin sites. To assess genome wide MyoD1 binding, we performed CUT&RUN and sequencing of Halo-MyoD1 in our C2C12^MyoD1::Halo + iMyoD1^ cell line treated ±dox for 6 h, a time point corresponding to maximum dox-induced MyoD1 dosage **(Fig. 1D, Fig. 3C)**. We recovered 9350 reproducible MyoD1 CUT&RUN peaks without treatment, and 13712 peaks with dox treatment. Motif enrichment analysis of the no-dox peak set recovered the MyoD1 E-box motif and the motifs of MyoD1 homologs and heterodimer partners, as has been previously reported for MyoD1-ChIP **(Fig. S5A)**.^17^ We also confirmed that Halo-MyoD1 bound to the previously reported target genes *Mef2a, Id3, MyoD1* and *Mymk* **(Fig. 3E, S7A-D)**.^17^ The +dox consensus peak set largely overlapped the untreated peak set and substantially broadened beyond it to recover 5310 new peaks, which we termed as “expanded” targets **(Fig. 3D)**.

To assess MyoD1-dose-dependent binding preference, we analyzed ±dox CUT&RUN signal intensity at MyoD1 “core targets” (peaks recovered at endogenous MyoD1 levels) or “expanded targets” (peaks recovered at elevated MyoD1 dosage) **(Fig. 3D)**. We found that dox treatment increased MyoD1 binding at both core and expanded targets **(Fig. 3F)**, with core targets being generally more highly occupied at basal MyoD1 expression levels than expanded targets **(Fig. S5E)**. Notably, comparing relative binding preference, we saw that dox treatment preferentially increased binding at expanded targets (1.93-fold) vs. core targets (1.57-fold) **(Fig. 3F)**. This result is reflected by some increase in binding at the core target *Mef2a*, and comparatively more dox-related binding at the expanded target *Cdh23* **(Fig. 3E)**. Together, these findings suggest that expanded MyoD1 targets are otherwise lowly occupied sites, whose binding is preferentially increased at elevated MyoD1 dosage.

MyoD1 has been reported to exhibit pioneer factor-like properties during myogenic reprogramming,^33^ so we sought to test whether elevated MyoD1 levels induced widespread chromatin remodeling to account for expanded genome-wide MyoD1 binding. We performed ATAC-sequencing of C2C12^iMyoD1^ cells treated with either 24 h of dox or vehicle, which produced modest changes in chromatin accessibility in both proliferating and differentiating myoblasts (proliferating: 43 regions gained and 23 lost; differentiating: 276 gained, 365 lost) **(Fig. S6A-D)**. These effects were far smaller than the accessibility changes due to 24 h of endogenous myogenic differentiation (1838 gained, 1455 lost) **(Fig. S6A-D)**. This limited effect may reflect the pre-existing myogenic regulatory environment in C2C12^iMyoD1^ cells, where endogenous MyoD1 and related muscle regulatory factors have already established much of the accessible chromatin landscape. In contrast, prior studies reporting MyoD1-dependent chromatin remodeling during transdifferentiation examined longer time windows over larger cell-state transitions, which may have produced more extensive accessibility changes.^34^ Next, we sought to test the extent of MyoD1 binding expansion occurring via binding of already accessible chromatin. We intersected the expanded MyoD1 CUT&RUN peaks with ATAC-seq peaks from untreated proliferating myoblasts or ATAC peaks unique to dox treatment. We found that 84% of expanded MyoD1 binding occurred in chromatin that was already accessible in untreated myoblasts, while only 7% of expanded MyoD1 binding occurred at dox-unique ATAC peaks **(Fig. S6E)**, in agreement with expanded MyoD1 targets being somewhat occupied even at basal MyoD1 expression levels **(Fig. S5E)**. Together, these results suggest that the observed expansion in MyoD1 binding occurs at generally accessible chromatin.

We next wondered if elevated MyoD1 expression expanded binding via the engagement of lower-affinity motifs. Indeed, we found that fewer expanded MyoD1 binding site peaks contained the canonical “CAGCTG” sequence **(Fig. 3G)**. Position-weight-matrix (PWM) analysis confirmed that expanded MyoD1 peaks scored lower for the consensus MyoD1 PWM **(Fig. S5D)**. These findings suggest that increased MyoD1 dosage expands MyoD1 binding to otherwise less frequently occupied, lower-affinity sites. We next performed motif enrichment analysis of expanded MyoD1 peaks to identify potential new binding motifs, which enriched TGAC/GTCA motifs related to Fra/Fos/Jun (AP-1) factors, and MyoD1-like E-box motifs at a lower rank order **(Fig. S5B**,**C)**. Because MyoD1 has been reported to interact with Jun family proteins, one possibility is that these motifs reflect cooperative binding. Alternatively, these motifs could reflect accessible chromatin featuring high AP-1 activity. Motif enrichment of ATAC-seq peaks in proliferating myoblasts (without dox) also recovered TGAC/GTCA motifs, suggesting that expanded MyoD1 binding occurs alongside motifs enriched within generally accessible myoblast chromatin **(Fig. S6F)**. TOBIAS footprinting analysis of ATAC peaks that overlapped with core and expanded CUT&RUN binding revealed that MyoD1 induction increased footprinting primarily via myogenic bHLH/E-box motifs, including that of known heterodimer partner Tcf12. These results suggest that elevated MyoD1 dosage enables the binding of lower-affinity E-box-like-motifs in accessible chromatin, likely in combination with known MyoD1 heterodimer partners such as Tcf12, rather than by directly engaging novel interaction partners such as AP-1 **(Fig. S6G-H)**.

To probe the gene regulatory effects of expanded MyoD1 binding, we correlated expanded MyoD1 binding to RNA sequencing of C2C12^iMyoD1^ cultures. Heatmap representation demonstrated a positive trend between expanded MyoD1 binding and gene upregulation **(Fig. 3H)**. By sorting peaks based on MyoD1 consensus motif presence, we observed that a trend towards upregulation was associated with expanded binding regardless of peak “CAGCTG” consensus sequence presence **(Fig. 3H)**.

Our integration of SMT, CUT&RUN and RNA-sequencing results suggest that increased MyoD1 dosage broadened genome-wide binding via engagement of lower-affinity sites. Therefore, we propose that increasing MyoD1 dosage can expand its gene regulatory repertoire by engaging lower-affinity regulatory elements in a manner compatible with mass-action binding. We term this mode “gene regulatory spillover”, whereby otherwise lowly expressed and lowly MyoD1-occupied genes are upregulated by additional chromatin engagement. Next, we sought to investigate whether dose-responsive MyoD1 targets could recapitulate the contractile phenotype.

### MyoD1 dose-responsive cell adhesion genes promote myotube contraction

To identify potential target genes linking elevated MyoD1 dosage to the contraction phenotype, we performed gene ontology analysis of MyoD1-bound and dose-responsive genes. Gene ontology analysis revealed an enrichment of cell junction and cell peripheral processes **(Fig. 4 A)**. Previous reports have suggested that wild-type C2C12 myotubes are capable of spontaneous contraction, but often detach and hyper-contract thereafter.^22^ We therefore hypothesized that the MyoD1-dose responsive program may enhance adhesion of the differentiated cell population to prevent detachment and thereby enable contraction. Consistent with this model, differentiated, dox-treated C2C12^iMyoD1^ were slower to detach in response to trypsin treatment as compared to untreated C2C12^iMyoD1^ **(Fig. S8A**,**B)**, suggesting enhanced adhesion of the whole differentiated cell population.

**Figure 4.**
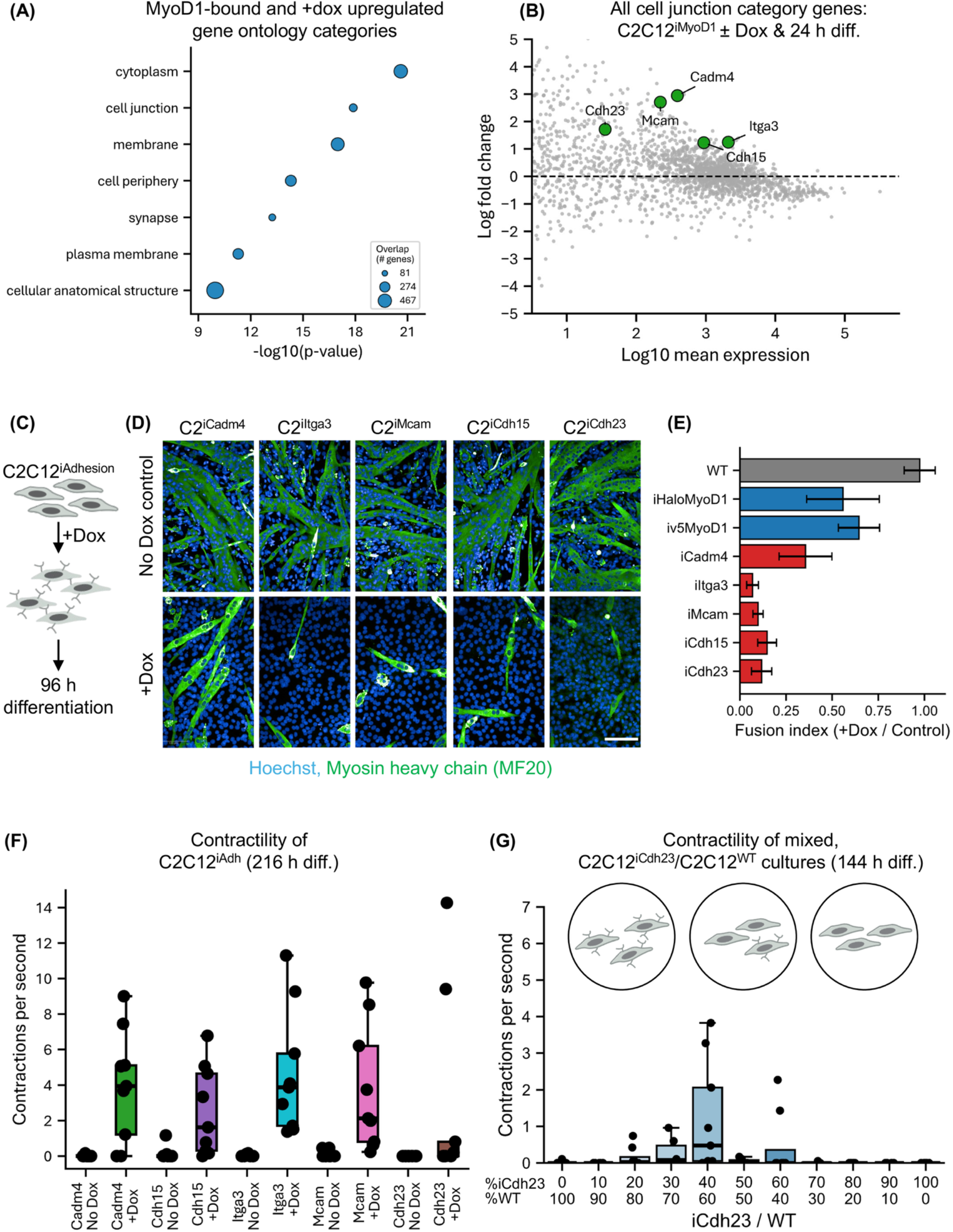
MyoD1-dose-responsive cell-adhesion genes enable myotube contraction. (A) Cell compartment gene ontology analysis of CUT&RUN MyoD1-bound genes that are significantly (adj. p <0.05) upregulated (2-fold) in +dox, 24 h diff. C2C12^iMyoD1^ RNA-seq. (B) MA plot of RNA expression of cell junction gene ontology category genes in C2C12^iMyoD1^ at 24 h diff. +dox. MA plot represents 3 independent RNA-seq biological replicates. Green dots annotate the expression of selected cell-cell adhesion factors *Cdh23, Cadm4, Itga3, Mcam*, and *Cdh15*. (C) Schematic describing the dox induction and 96 h differentiation of C2C12 bearing dox inducible adhesion genes (C2C12^iAdhesion^). Grey cells (top) represent uninduced myoblasts. Grey cells with semi-circle arms (middle) represent myoblasts expressing adhesion molecules. (D) Representative confocal microscopy images of Myosin heavy chain (MF20, green) immunoreactivity and Hoechst staining (blue) of C2C12 cell lines treated ± dox simultaneous with the onset of 96 h differentiation. Columns represent C2C12^iCadm4^, C2C12^iItga3^, C2C12^iMcam^, C2C12^iCdh15^ and C2C12^iCdh23^ cell lines. Scale bar: 100 µm. (E) Quantification of (D) representing the percentage of nuclei fused into MyHC immunoreactive myotubes at 96 h differentiation. Fusion index represents the percentage of fused nuclei in +dox condition divided by no dox control. Analysis represents measurements of >100,000 nuclei over 3 biological replicates. Bars represent fusion index of C2C12^WT^, C2C12^iHaloMyoD1^, C2C12^iV5MyoD1^, C2C12^iCadm4^, C2C12^iItga3^, C2C12^iMcam^, C2C12^iCdh15^ and C2C12^iCdh23^. Error bars represent SEM of biological replicates. (F) Optical flow contraction analysis of C2C12^iCadm4^, C2C12^iCdh15^, C2C12^iCdh23^, C2C12^iItga3^ and C2C12^iMcam^ cultures treated ±dox and differentiated 216 h. Dots represent 9 well replicates across 3 biological replicates; line represents median of well replicates, box edges represent quartile 1 (lower) and quartile 3 (upper), whiskers represent 1.5× IQR. (G) Optical flow contraction analysis of mixed C2C12^iCdh23^/C2C12^WT^ cultures treated +dox and differentiated 144 h. Dots represent 4-9 well replicates across 2-3 biological replicates. First row of x-labels describes percentage of cell population that is C2C12^iCdh23^, second row describes percentage of population that is C2C12^WT^.

We next selected candidate adhesion-promoting genes that were both dox-upregulated and more MyoD1-bound, identifying *Cadm4, Itga3, Mcam, Cdh15* and *Cdh23*, all previously reported to mediate cell adhesion **(Fig. 4B, Fig. S7E-H)**.^35^ To test whether induction of individual candidates was sufficient to recapitulate the contraction phenotype, we generated C2C12 harboring dox-inducible *Cadm4, Itga3, Mcam, Cdh15*, or *Cdh23* alleles (collectively C2C12^iAdhesion^). Ectopic expression of each adhesion gene inhibited myogenic differentiation to a greater extent than the induction of Halo-MyoD1 or V5-MyoD1 **(Fig. 4C-E)**. Nevertheless, we also observed contracting myotubes in most replicates of *Cadm4, Itga3, Mcam* and *Cdh15*-expressing cultures after 216 h of differentiation, 72 h later than the spontaneous contraction observed in C2C12^iMyoD1^ **(Fig. 4F, Supplementary Video 2)**. Stronger differentiation inhibition due to adhesion gene expression may have delayed myoblast fusion and/or myotube maturation and thus the onset of spontaneous contraction.

The observation that both elevated MyoD1 levels, and the ectopic expression of individual adhesion genes inhibited fusion suggested a two-population mechanism, in which enhanced adhesion in the unfused part of the population could anchor and stabilize the contraction of myotubes formed by more fusion-competent cells. To test this potential mechanism, we differentiated mixtures of differentiation-competent myoblasts (C2C12^WT^) with myoblasts expressing *Cdh23* (C2C12^iCdh23^) that were differentiation-inhibited. Neither of these cell populations formed contractile myotubes when differentiated alone for 144 h **(Fig. 4G)**. Cell mixtures containing between 30-60% C2C12^iCdh23^ / 70-40% C2C12^WT^ formed contracting myotubes after 144 h of differentiation and dox treatment, indicating that *Cdh23* expression in a subset of the cell population supported contraction of myotubes likely formed primarily by wild-type C2C12 **(Fig. 4G)**. We next asked whether a sub-fraction of C2C12^iMyoD1^ myoblasts could support myotube contraction. Cell mixtures containing between 50-100% C2C12^iMyoD1^/C2C12^WT^ featured contracting myotubes, where cultures containing 30-40% C2C12^WT^ most reliably formed contracting myotubes **(Fig. S8C)**. This trend suggests that increasing the fraction of more differentiation-competent cells enhances formation of myotubes, where a MyoD1-high subpopulation may provide a complementary adhesive contribution to stabilize myotube contraction.

Next, we asked whether the MyoD1 dose-responsive adhesion program was preferentially associated with unfused myoblasts at later stages of C2C12^iMyoD1^ differentiation. To address this, we differentiated C2C12^iMyoD1^ ±dox for 120 h and isolated myoblast-enriched and myotube-enriched fractions via trypsinization and filtration **(Fig. S9A)**.^36^ This strategy yielded fractions with expected transcriptional identities, where myoblast fractions expressed higher levels of proliferation-associated markers, whereas the myotube fractions showed higher expression of myosins, troponins, titins, and other myotube-associated genes **(Fig. S9B)**. Dox treatment produced a transcriptional response in the myoblast-enriched fraction, but little significant response in the myotube-enriched fraction (myoblast: 252 upregulated, 452 down, myotube: 4 up, 2 down) **(Fig. S9C**,**D)**. Replicate variability limited significance of gene regulatory changes in the myotube-enriched ±dox comparison. The dox-upregulated genes in the myoblast fraction included adhesion-related candidates *Cadm4* and *Mcam*, whereas none of the candidate adhesion genes were upregulated in the myotube fraction. Consistent with this pattern, gene ontology analysis of genes upregulated in the myoblast-enriched fraction recovered cell projection and cell periphery terms **(Fig. S9E)**, while the myotube-enriched fraction did not yield significantly enriched ontology terms. Together, these results indicate that the MyoD1 dose-responsive adhesion program is most clearly detected in the unfused myoblast-enriched population at 120 h.

To independently assess how the adhesion gene program contributed to the MyoD1 dose-responsive transcriptional state, we also performed RNA-seq of C2C12^iMcam^ and C2C12^iCdh23^ lines treated ±dox and differentiated for 48 h **(Fig. S10A)**. Induction of either Mcam/Cdh23 upregulated additional adhesion-associated genes, along with broad downregulation of many genes related to myogenic differentiation, including fusogens, myogenic regulators, and sarcomere genes **(Fig. S10B)**. Cdh23- and Mcam-upregulated genes overlapped substantially with the MyoD1-induced program, encompassing ∼70% of MyoD1-upregulated genes at 48 h of dox treatment and differentiation **(Fig. S10C)**. Gene-wise fold-change comparisons showed that Cdh23 and Mcam induction correlated with MyoD1 induction, and most strongly with each other (Spearman ρ = 0.62 (Cdh23 vs. MyoD1), 0.67 (Mcam vs. MyoD1) and 0.81 (Cdh23 vs. Mcam)) **(Fig. S10D-F)**. Finally, the mean Cdh23/Mcam-induced response was more strongly correlated with the MyoD1-induced response in the myoblast-enriched fraction than in the myotube-enriched fraction **(Fig. S10G)**. Together, these results suggest that MyoD1-responsive adhesion genes reinforce an adhesion-associated myoblast state rather than primarily altering myotubes. Secondly, these results imply that many of the downstream transcriptomic aspects of early elevated MyoD1 levels can be mediated through an adhesion axis.

Taken together, these results are compatible with a model in which elevated MyoD1 levels, or the induced expression of adhesion-promoting genes, generated an adhesive population that supported the contraction of myotubes formed by fusion-competent myoblasts. Because multiple individual adhesion genes were sufficient to impair fusion and promote contraction, we suggest that this phenotype is unlikely to depend on a single uniquely required adhesion factor but instead reflects a partially redundant adhesion program activated by elevated MyoD1 dosage.

## Discussion

In this study, we employed an inducible expression system to test the relationship between elevated MyoD1 expression and the gene regulation of myogenic differentiation. We report that, contrary to our original expectations, increased MyoD1 dosage inhibited multiple molecular and phenotypic hallmarks of canonical myogenesis while enabling a robust myotube contraction phenotype. Increased MyoD1 dosage did not further amplify genes normally upregulated by myogenic differentiation but instead upregulated normally lowly expressed genes. Single-molecule tracking and CUT&RUN analysis revealed that MyoD1 dosage regulated the total amount of MyoD1 molecules engaging chromatin, where excess MyoD1 preferentially engaged lower-affinity sites in a manner compatible with mass-action binding predictions. Expanded engagement correlated with the upregulation of nearby genes. We found that the ectopic expression of select MyoD1 dose-responsive adhesion genes similarly inhibited myogenic differentiation and enabled myotube contraction. Together, these results suggest that MyoD1 dosage regulates an ensemble of genes via chromatin binding over an affinity gradient.

The MyoD1-dose responsive “off-script” adhesion gene program may explain the superficially paradoxical phenotype of myotube contractility despite myoblast fusion inhibition. Cell behaviors required for fusion, such as migration, alignment, and membrane remodeling may be inhibited by increased myoblast adhesion.^37^ Secondly, elevated MyoD1 dosage may have also altered the stoichiometry of available co-factors and potentially titrated MyoD1 E-protein heterodimer partners away from canonical myogenic loci towards expanded binding sites, contributing to the observed downregulation of the endogenous myogenic program. We suggest that the contractile phenotype arose through elevated MyoD1 (or single adhesion-gene induction) increasing the fraction of unfused, and adhesive myoblasts that behave like an “adhesive scaffold,” stabilizing developing myotubes against detachment to enable sustained contractility. Importantly, our mixing experiments show that an adhesion-gene-high subpopulation is sufficient to support contraction in mixed cultures containing otherwise differentiation-competent C2C12^WT^ cells, supporting a non-cell-autonomous mechanism **(Fig. 4G**). While the mixing experiments do not fully exclude heterotypic fusion into myotubes, our fractionated RNA-seq experiments corroborate that MyoD1 induction preferentially upregulates adhesion genes in unfused myoblasts **(Fig. S9)**.

We observed that both core and expanded MyoD1 targets were more highly bound at dox-induced MyoD1 dosages **(Fig. S5E)**, indicating that core sites were not fully saturated at dox-induced MyoD1 levels. Instead, these results suggest that MyoD1 binding increased according to mass-action by preferentially engaging lower-affinity target sites in accessible chromatin **(Fig. S6E)**. The observation that normally lowly expressed targets were preferentially responsive to increased MyoD1 dosage **(Fig. 2E)** is compatible with a gene activation threshold overcome by increased MyoD1 binding. Therefore, we propose a mechanism of “gene regulatory spillover”, in which increased TF dosage enables broader genome-wide binding over an expanded binding site affinity gradient. Subsequently, broader binding may increase the likelihood of transcriptionally activating normally unresponsive genes, thereby inducing an alternate transcriptional program that alters cellular decision-making.

Dose-dependent engagement of lower-affinity regulatory elements has also been described for other lineage-specific TFs capable of cell reprogramming. For example, Oct4, Sox2, Klf4 and c-Myc dosage have been reported to control the binding of low-affinity sites during the pluripotency reprogramming of fibroblasts.^38^ Investigation of dose-dependent c-Myc function has found the “invasion” of lower-affinity cis-regulatory elements.^39,40^ We similarly found that MyoD1 dose-dependent binding expansion occurred in constitutively accessible chromatin **(Fig S6E)**. Therefore, we speculate that gene regulatory spillover may emerge from both the biophysical principles governing TF binding and the on/off switch nature of gene activation. Given orthogonal examples of TF dose-dependent cell fate determination, we propose that gene regulatory spillover may be a viable mechanism underpinning the role of multiple TFs in directing cell state changes.

## Materials and Methods

### Cell culture

Murine C2C12 myoblasts (ATCC CRL-1772) were maintained at 37 °C in a humidified 5% CO_2_ atmosphere in “Growth medium” (GM) containing DMEM (4.5 g/L glucose) supplemented with 20% fetal bovine serum (FBS), 1X penicillin–streptomycin, GlutaMAX and sodium pyruvate. Cells were passaged every 3–4 days at ∼80% confluence via 1X PBS wash, trypsinization employing 0.05% trypsin and re-seeding at a 1:10 ratio. Cells were trypsinized and counted using a Countess automated cell counter prior to seeding for all experiments. C2C12 cultures were confirmed negative for mycoplasma by in-house PCR and agarose-gel electrophoresis.

### Myogenic differentiation

C2C12 myoblasts were grown to 90% confluence in GM. Differentiation was induced by switching myoblasts to “Differentiation medium” (DM), composed of DMEM containing 2% horse serum, GlutaMAX, 1X penicillin–streptomycin, sodium pyruvate and 1X ITS supplement (insulin, transferrin and selenium). Differentiation was performed without medium replacement for up to 216 h.

### Transfection, stable cell line generation and dox induction

2×10^5^ C2C12 cells were plated in plastic 6-well culture dishes containing GM media and allowed to proliferate for 24 h prior to transfection. C2C12 were transfected with 5 µg total plasmid DNA (4 µg dox inducible XLone transgene vector + 1 µg Super PiggyBac transposase) using Lipofectamine 3000 following manufacturer protocols. Media was replaced 24 hours post-transfection with GM containing Puromycin. Stably integrated cells were selected using 1 µg/mL Puromycin. XLone-based cell lines were induced to express constructs via the addition of 1 µg/mL doxycycline.

### Plasmids employed to generate dox-inducible cell lines

XLone TRE3G 3xF-Halo-GDGAGLIN-Myod1

XLone TRE3G V5-Halo-GDGAGLIN-Myod1

XLone TRE3G V5-Myod1

XLone TRE3G Halo-NLS

XLone TRE3G Cdh23

XLone TRE3G Cadm4

XLone TRE3G Itga3

XLone TRE3G Cdh15

XLone TRE3G Mcam

Super PiggyBac Transposase (PB200PA-1)

### Immunohistochemistry

2×10^4^ C2C12 cells were seeded in 96-well Perkin Elmer PhenoView dishes containing GM. Cells were grown to ∼90% confluence and induced to differentiate as described above. For Halo-MyoD1 visualization, cultures were stained with 50nM of JFX549 HaloTag ligand for 15 minutes, washed 2x with phenol free DMEM 20% FBS and destained for 15 minutes in DMEM 20% FBS. Cells were fixed with 4% paraformaldehyde diluted in PBS for 10 min at 37 deg. C, permeabilized and antigen blocking was performed in PBS containing 0.25% Triton-X100 and 2% bovine serum albumin (BSA) for 1 h at room temperature (RT), incubated with primary antibody in PBS containing 0.125% Triton-X100 and 2% bovine serum albumin at 4°C overnight, washed 3x with PBS, then incubated with secondary antibody in PBS containing 0.25% Triton-X100 and 2% bovine serum albumin at RT for 1 hr. Post-antibody staining, C2C12 nuclei were labeled with Hoechst 33342 at 1:10,000 in PBS for 10 minutes, washed 3x with PBS, and imaged in PBS.

### Confocal microscopy

Confocal microscopy was performed on a Perkin Elmer Opera Phenix microscope. For all confocal microscopy experiments, a 20x water NA=1.0 objective was used to acquire up to 3 channels, covering >30% each well per replicate.

The following microscopy parameters were employed for confocal imaging: 375/435–480 nm (Hoechst, 40 ms, 100% laser), 488/500–550 nm (Myogenin, 100 ms, 50%; MF20, 500 ms, 100%), and 561/570–630 nm (Halo-JFX549, 500 ms, 100%). For each experiment, Z-stacks covering all nuclear planes were acquired. Multiple fields of view were acquired to cover >30% around the center of each well per technical well replicate.

### Confocal microscopy analysis

All confocal microscopy analysis was performed employing Perkin Elmer Harmony software. *Nuclear segmentation:* Maximum intensity projection and basic flatfield correction was applied to all image z-stacks. Nuclei were segmented on Hoechst labeling using the “Find Nuclei” module, Method M, applying a 30 µm diameter, 0.4 splitting sensitivity and 0.00 common threshold.

*Myotube segmentation:* Myotubes were segmented on MF20 immunohistochemistry intensity employing the “Find Cells” module, Method C, Common Threshold 0.2, Area>50 µm^2^. Cells were clustered via “Cluster by Distance method” (1px, Area>10000 pix.) to join over-segmented myotubes. *Myotube fused nuclei:* To determine the percentage of nuclei fused into myotubes, we masked nuclei by Myotube segmentation-generated masks. Overlap of nuclei to myotube mask was determined by geometrical center of nuclei to myotube mask positioning. Percent myoblast fusion was calculated by taking the ratio of the number of fused to all nuclei (Fused nuclei/All nuclei)*100. *MyoD1/MyoG-expressing nuclei:* To identify nuclei expressing MyoD1 and MyoG, nuclei were segmented on Hoechst intensity as described above. Nuclei were called as MyoD1/MyoG expressing if their fluorescence intensity reached above a threshold defined by negative control (2ndry antibody only).

### Western blotting

C2C12 (1 × 10^6^) were harvested by direct RIPA buffer (CSHL) + 25U Benzonase lysis. Lysates were mixed with 4× SDS-PAGE loading buffer (CSHL) to a final 1× concentration and run on 4– 12% Bis-Tris polyacrylamide gels at 100V for 1.5 h. Proteins were transferred to 0.45 µm nitrocellulose membranes overnight at 10V, 4 °C in transfer buffer containing 20% methanol. Membranes were blocked in TBS with 0.2% Tween-20 and 3% BSA at room temperature for 1 h. Blocked membranes were incubated with primary antibodies (anti-MyoD1, anti-TBP, anti-FLAG) overnight at 4 °C. After primary antibody probing, membranes were washed 3x in TBS with 0.2% Tween-20 and probed with secondary HRP-conjugated antibody diluted in TBS with 0.2% Tween-20 and 3% BSA. Blots were either cut and separately probed or serially probed for TBP detection. Detection was performed using HRP-conjugated secondary antibodies and chemiluminescent substrate (WestECL) and imaged on a Bio-Rad Chemidoc system. Western blots were quantified in Fiji (ImageJ).

### Antibodies

Anti-MYH1E – MF 20, (Developmental Studies Hybridoma Bank), 2 µg/mL; Anti-MyoG – ab1835, (Abcam), 2 µg/mL; Anti-IgG (H+L) Alexa Fluor 488 – A11001, (Molecular Probes), 2 µg/mL; Anti-MyoD1 - MA5-12902, (Thermo Fisher), 0.2 µg/mL; Anti-TBP – mAbcam51841, (Abcam), 1 µg/mL; Anti-Light chain HRP – Jackson ImmunoResearch, 0.08 µg/mL; Anti-flag M2– F3165, (Sigma-Aldrich), 4.7 µg/mL.

### Contraction assay and phase-contrast microscopy

C2C12 lines were seeded at 150,000 cells per well of a 24-well culture dish, grown to 90% confluence and differentiated for 144-216 h as described above. Phase-contrast microscopy was performed using a 4x air NA=0.1 Ph0 objective or 10x air NA=0.25 Ph1 objective. One representative field of view featuring myotubes was recorded over well replicates; all analyses were performed across 3 biological replicates per condition unless otherwise indicated.

### Trypsinization assay

C2C12 myoblasts were differentiated for 120 h to assess adherence prior to the onset of contraction. Cells were washed once with 2 mL room-temperature PBS and temperature equilibrated for 10 min in 0.5 mL PBS. Detachment was induced by adding 2 mL of 0.05% trypsin at the start of brightfield time-lapse imaging, and each condition was recorded for 5 min. The time of myotube detachment onset was manually scored for each well.

### Optical flow contraction analysis

Contraction analysis was performed by implementing OpenCV2-computervision to quantify whole-time-lapse cellular motion using dense Farneback optical flow. Videos (∼15 -30 s duration) were first trimmed by 5 s at both start and end to exclude artifacts. Pairwise optical flow was computed with OpenCV’s calcOpticalFlowFarneback (pyr_scale = 0.5, levels = 3, winsize = 15, iterations = 3, poly_n = 5, poly_sigma = 1.2, flags = 0). The resulting horizontal (u) and vertical (v) flow vectors were converted to magnitude and summed over all pixels to yield a “motion” scalar per interframe interval. For motion over time visualization, values were smoothed using a centered moving average (2-frame) window. For contraction-rate plots, motion-scalar timepoints within a predefined motion range were counted and divided by the analyzed video duration to yield contractions per second.

### Cas9 Halo tagging

Endogenous Halo-tagging was performed as previously described.^41^ C2C12 were plated at 500,000 cells per 10cm culture dish, grown to 40% confluence and transfected as described above with Cas9 editing reagents consisting of a pENTR-based N-terminally Halo-tagged MyoD1 homology-dependent repair donor plasmid and pU6 sgRNA, Cas9-Venus-bearing editing plasmid.

Guide RNA sequence for MyoD1 halo-tag knock-in:

pU6_gRNA144_myod: 5’ gagcttctatcgccgccactc 3’

Homology-dependent repair (HDR) vectors included 200-500 bp long arms homologous to the *MyoD1* genomic region.

HDR and gRNA plasmids:

pHR_mMyoD_NtermHaloKI_gRNA144mut

pU6_gRNA144_myod

Post-transfection, cells were cultured for 24 h, media was exchanged, cells were cultured for another 24 h and then FACS-sorted into 96-well plates based on Cas9-Venus fluorescence intensity to select for transfected cells. Single-cell clones were grown for an additional 2 weeks and then genotyped for inclusion of N-terminal MyoD1 Halo-tag via genomic PCR screening, Sanger sequencing and western blotting.

### Single-molecule imaging experiments

For single-molecule imaging experiments, C2C12 cells were grown on glass-bottom 35mm MatTek dishes and were stained with 50nM of JFX549 Halo-tag ligand for 15 minutes, destained as for confocal microscopy, and imaged within 4 h of Halo-tag labeling.

Single-molecule microscopy was performed as previously detailed, employing a Nikon TI microscope with a 100X/NA 1.49 oil objective.^42,43^ Time-lapses were acquired employing a 1 watt 561 nm laser. For Halo-MyoD1 microscopy, an initial snapshot of cells was acquired, after which cells were bleached at maximum laser power in 150×150 ROIs for 10-30 seconds to generate single-molecule labeling density of Halo-MyoD1. Time-lapses were acquired within 150×150 pixel ROI manually applied to the center of nuclei. Time-lapses were captured with stroboscopic illumination at 1 msec exposure at 561nm excitation. Five to ten thousand frames were recorded per cell, depending on experiment labeling density. A minimum of 20 cells were recorded per condition, with 2 independent biological replicates recorded on separate days and pooled for analysis.

### Single-molecule detection, localization and tracking

Detection, localization and tracking of molecules in single-molecule movies were processed as previously detailed.^44^ In brief, we employed the ‘quot’ package (https://github.com/alecheckert/quot), using the following tracking parameters: method = ‘conservative’, pixel_size_um = 0.160, frame_interval = 0.00748, search_radius = 1.0, max_blinks = 0, min_I0 = 0.0, scale = 7.0.

### Nuclear masking of single-molecule trajectories

Trajectories were masked by nuclear snapshots to minimize analysis contamination by non-nuclear trajectories. In brief, saturation HaloTag ligand labeled nuclear snapshot images of Halo-MyoD1 were Gaussian blurred and thresholded to produce a binary mask. Hole filling was applied to this binary mask. Mask contours were extracted and applied to single-particle trajectory localizations.

### Inference of diffusivity from single-molecule tracking

Diffusivity inference of single-molecule trajectories was performed with a Bayesian state array approach as previously detailed.^44^ In brief, we employed the “saspt” package (https://github.com/alecheckert/saspt).^44^ To discard time-lapse start artifacts, the first 1000 frames of each set of tracks were excluded from analysis. Chromatin bound fractions were extracted from saspt state arrays via thresholding at 0.1 µm^2^/s to define fraction chromatin bound.

### MyoD1 fluorescence intensity analysis

Snapshots of saturation-labeled Halo-tag ligand staining were normalized and gaussian blurred prior to use as nuclear segmentation masks. Nuclear segmentation was performed via inverse Otsu thresholding and applied to the original snapshot to compute mean fluorescence intensity.

### RNA-sequencing and analysis

C2C12 myoblasts were seeded at 200,000 (differentiation condition) or 100,000 cells (undifferentiated condition) per 6-well dish, treated with 1 µg/mL doxycycline or vehicle, and simultaneously induced to differentiation or maintained in growth media. Total RNA was extracted using TriZol following manufacturer protocols. polyA-mRNA enrichment was performed on extracted RNA using the NEBNext Poly(A) mRNA Magnetic Isolation Module (E7490S). Extracted RNA was reverse transcribed following NEBNext Ultra II RNA Library Prep kit, following manufacturer protocol. Libraries were prepared following NEBNext Ultra II RNA Library Prep kit protocol. Sequencing was performed by MedGenome (Foster City, CA) on Illumina NovaSeq/NovaSeq 6000 instruments with 150bp paired end reads.

Quality control and processing was performed with the nf-core/RNAseq pipeline version 3.14.0.^45^ Differential expression testing was performed with DESeq2 in R (design ∼ condition), with gene annotation via biomaRt. Gene ontology enrichment was performed with gProfiler. MA plots were generated in Python with Matplotlib, pandas, and NumPy. Heatmaps of selected targets were drawn using pheatmap and RColorBrewer in R, while three-way Venn diagrams of upregulated genes were generated using matplotlib-venn.

RNA-seq. analyses reflect 3 biological replicates per condition and time point, except for the +dox-treated myoblast-enriched condition, which contains 2 biological replicates.

### Myoblast/Myotube filtration

To separate differentiated cultures into mononucleated and myotube-enriched fractions, cells were treated with 0.00125% trypsin for 5 minutes, triturated ten times, and passed through a 40 µm cell strainer. The flow-through was collected as the mononucleated myoblast-enriched fraction, while the retentate on the filter was collected as the myotube-enriched population. The retentate was collected by backflushing the filter using PBS. Both flow-through and retentate were centrifuged at 500xg for 5 min and lysed in TriZol prior to RNA extraction.

### CUT&RUN sequencing and analysis

C2C12^MyoD1::Halo + iMyoD1^ and C2C12^WT^ were seeded at 200,000 cells in a 6-well dish, grown to ∼90% confluence and induced to differentiation simultaneous with +/-1 µg/mL doxycycline treatment for 6 hours. CUT&RUN was performed according to published protocols.^46^ In brief, 500,000 cells were harvested via trypsinization, permeabilized, incubated with M2-anti-flag antibody and treated with pAG-MNase. Fragments were purified using phenol-chloroform DNA extraction. Library preparation was performed using NEBNext Ultra II DNA library Prep kit for Illumina, following manufacturer protocols. Sequencing was performed by MedGenome (Foster city, CA) on an Illumina NovaSeq X instrument with 150 bp paired-end reads.

Fastq files were analyzed using the nf-core/cutandrun pipeline version 3.2.2.^45^ Pipeline parameters that deviated from default: genome: mm10, normalization_mode: CPM, replicate_threshold:2, seacr_stringent: relaxed, trim_nextseq: 20, blacklist:mm10-blacklist.v2. Motif enrichment analysis was performed using HOMER. Genome track visualization was performed using IGV 2.17.4. CUT&RUN binding intensity analysis was performed by scaling per-replicate BigWig coverage to counts-per-million, binning ±1 kb around each peak summit into 50 bp windows, averaging across replicates per condition, and plotting the resulting mean ± variability for each peak set. All CUT&RUN analyses reflect 3 biological replicates per condition.

### ATAC sequencing and analysis

C2C12^iMyoD1^ myoblasts were seeded at 300,000 (24 h differentiation condition) or 75,000 cells (undifferentiated condition) per well of a 12-well dish, treated with 1 µg/mL doxycycline, and simultaneously either induced to differentiation as described above (for 24 hours differentiated samples) or maintained in 20% FBS DMEM for 24 hours (undifferentiated samples). ATAC tagmentation and fragment purification was performed according to published protocols.^47^ Cells were harvested via trypsinization, and 20,000 were permeabilized and tagmented with 2 uL Illumina TDE1 enzyme. Fragments were purified using Zymo DNA clean and concentrator-5. Library preparation was performed using NEBNext Ultra II DNA library Prep kit for Illumina, employing custom Nextera primers. Sequencing was performed by MedGenome (Foster city, CA) on Illumina NovaSeq 6000 instruments with 100bp paired end reads.

Fastq files were analyzed using the nf-core/atacseq v 2.1.2 pipeline.^45^ Parameters that deviated from default: genome:mm10, blacklist: mm10-blacklist.v2, trim_nextseq:20, min_reps_consensus:2. All ATAC analyses reflect >=3 biological replicates per condition and time point.

TOBIAS analysis: ATAC-seq replicates were merged and analyzed with TOBIAS BINDetect separately for each peak class and comparison. Significance was assessed with a one-sample t-test against zero, requiring at least five sites per motif. TF-level volcano plots were generated from the mean effect size and p value. Motifs were first filtered to RNA-expressed TFs (baseMean ≥ 10), and differential hits were classified using the 7.5th/92.5th percentiles of effect size and the 92.5th percentile of -log10(p). ^48^

### CUT&RUN – ATAC - RNA-seq. multimodal analysis

Multimodal sequencing analyses were performed by merging ATAC and CUT&RUN consensus peak sets into a master consensus peak set and subsequently intersecting based on peak coordinates. Merged peaks were associated to the nearest annotated gene using HOMER. RNA-seq gene expression data were merged based on gene mapping via tx2gene.

### Use of AI

ChatGPT 5.5 was employed in preparing analysis code.

## Data availability

All sequencing data have been deposited to GEO and will be released upon publication.

## Acknowledgments

We thank Claudia Cattoglio for advice in carrying out CUT&RUN experiments.

We thank Djem Kissiov, Vinson Fan, Claudia Cattoglio and all members of the Tjian and Darzacq lab for critical readings of this manuscript.

We thank Mary West and the QB3 High-Throughput screening Facility (HTSF) at University of California, Berkeley. All confocal microscopy was performed in the QB3 HTSF (*RRID:SCR_022304*).

This work was supported by the Chan-Zuckerberg Initiative Dynamic imaging grant (to X.D.), and the Howard Hughes Medical institute (R.T.).

O.N.W was supported by the NIH training program grant T32GM007232. JKM was supported by the NIH training program grant T32GM139780.

