## Supplementary material for "Elevated MyoD1 levels expand genome-wide binding and the repertoire of regulated genes": SI Appendix

### FIG-S1

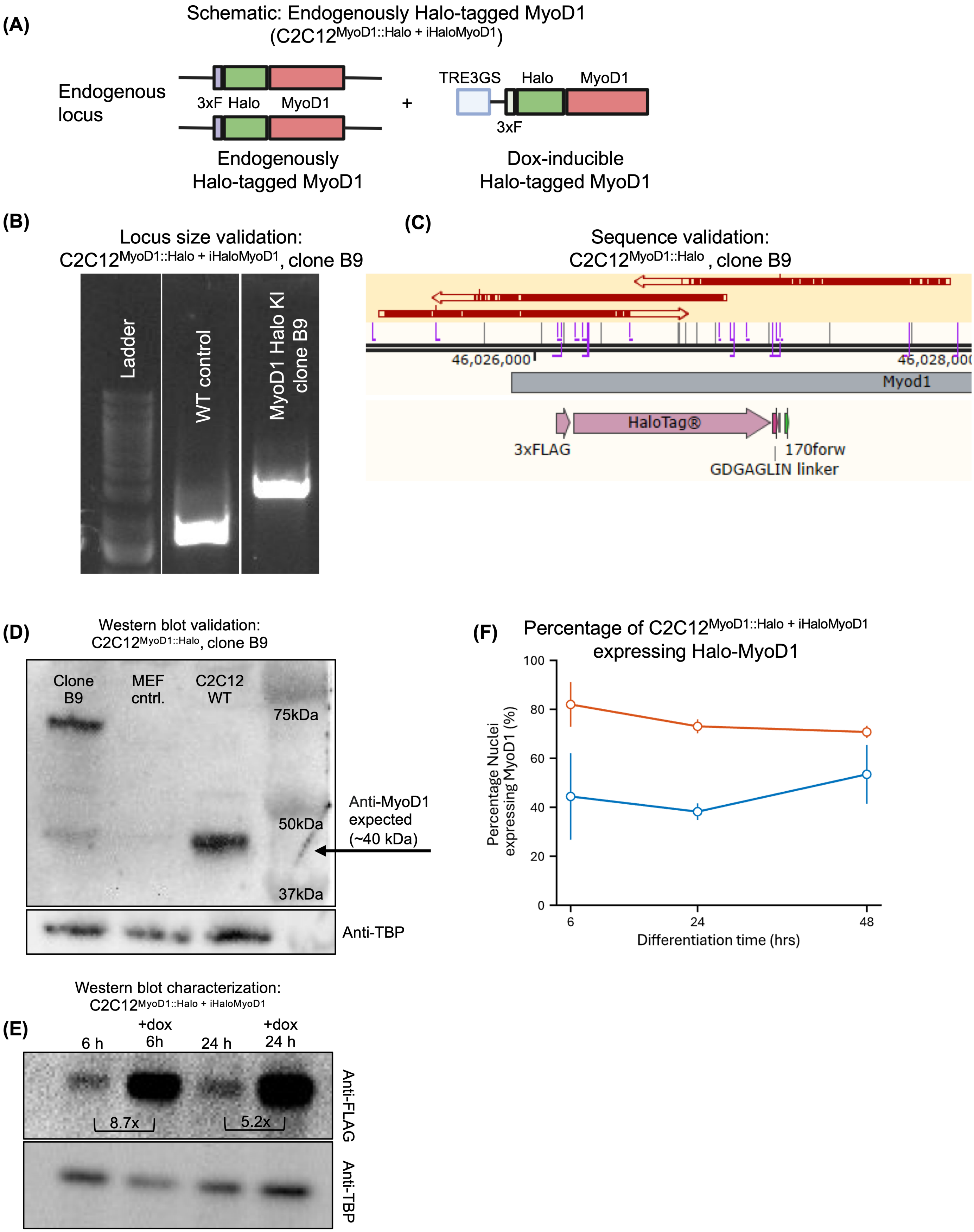

(A)
Schematic: Endogenously Halo-tagged MyoD1
(C2C12^MyoD1::Halo + iHaloMyoD1)
Endogenous locus
3xF Halo MyoD1
Endogenously Halo-tagged MyoD1
TRE3GS Halo MyoD1
3xF
Dox-inducible Halo-tagged MyoD1
(B)
Locus size validation: C2C12^MyoD1::Halo + iHaloMyoD1, clone B9
Ladder
WT control
MyoD1 Halo KI clone B9
(C)
Sequence validation: C2C12^MyoD1::Halo, clone B9
46,026,000
46,028,000
Myod1
HaloTag®
3xFLAG
170forw
GDGAGLIN linker
(D)
Western blot validation: C2C12^MyoD1::Halo, clone B9
Clone B9
MEF cntrl.
C2C12 WT
75kDa
50kDa
37kDa
Anti-MyoD1 expected (~40 kDa)
Anti-TBP
(F)
Percentage of C2C12^MyoD1::Halo + iHaloMyoD1 expressing Halo-MyoD1
Percentage Nuclei expressing MyoD1 (%)
100
80
60
40
20
0
6
24
48
Differentiation time (hrs)
Western blot characterization: C2C12^MyoD1::Halo + iHaloMyoD1
(E)
6 h
+dox 6h
24 h
+dox 24 h
8.7x
5.2x
Anti-FLAG
Anti-TBP

Supplementary Figure 1

(A) Schematic describing homozygous Halo-tagging of endogenous MyoD1 alleles and introduction of additional doxycycline-inducible Halo-MyoD1 alleles. Grey box represents 3x-Flag tag, green boxes represent HaloTag, red boxes represent *MyoD1*. White box represents dox-inducible *TRE3GS* promoter.

(B) PCR amplicons of MyoD1 N-terminal genome locus, indicating differential sizing of wild-type C2C12 (approx. 1600bp) and HaloTag knock-in clone (approx. 2600bp) MyoD1 N-terminal locus region.

(C) Sanger sequencing of homozygous N-terminal HaloTag knock-in clone B9, indicating Sanger sequencing coverage over the insertion of HaloTag sequence to MyoD1 N-terminus fusion. Arrows represent forward and reverse Sanger sequencing reads aligned to a reference *MyoD1* sequence. White dashes in arrows represent SNPs as compared to reference *MyoD1* sequence.

(D) Western blot probing MyoD1 in HaloTag knock-in C2C12$^{\text{MyoD1::Halo}}$ clone (B9), a mouse embryonic fibroblast negative control (Fibro. Control) and wild-type, parental C2C12 myoblasts (“WT”). Arrow marks expected wild-type anti-MyoD1 band size. Bottom lane indicates an anti-TBP control, spliced underneath the original anti-MyoD1 image.

(E) Western blot probing FLAG in HaloTag knock-in clone C2C12$^{\text{MyoD1::Halo + iMyoD1}}$, untreated or treated ±dox for 6 and 24 h. Brackets and annotation signify expression fold change normalized to TBP.

(F) Quantification of the percentage of C2C12$^{\text{MyoD1::Halo + iMyoD1}}$ cells expressing detectable Halo-MyoD1 over 48 h of differentiation and ±dox, as represented by (Fig. 1C). Orange line represents the fraction of +dox treated C2C12$^{\text{MyoD1::Halo + iMyoD1}}$, while blue line represents the fraction of untreated C2C12$^{\text{MyoD1::Halo + iMyoD1}}$. Plotted points and bars represent means and standard deviation across 3 biological replicates.

### FIG-S2

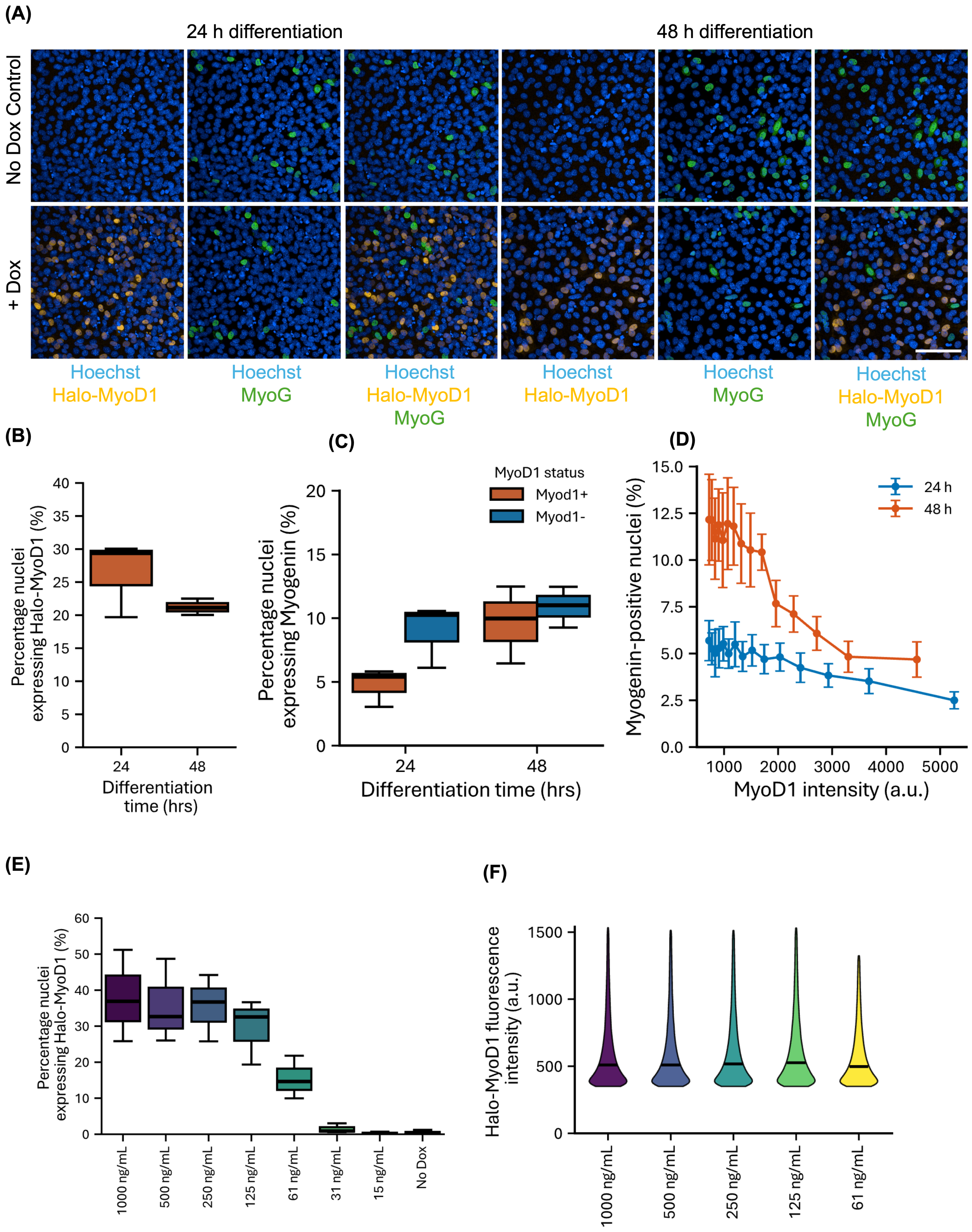

(A)
24 h differentiation
48 h differentiation
No Dox Control
+ Dox
Hoechst
Halo-MyoD1
Hoechst
MyoG
Hoechst
Halo-MyoD1
MyoG
Hoechst
Halo-MyoD1
Hoechst
MyoG
Hoechst
Halo-MyoD1
MyoG
(B)
Percentage nuclei expressing Halo-MyoD1 (%)
Differentiation time (hrs)
(C)
MyoD1 status
Myod1+
Myod1-
Percentage nuclei expressing Myogenin (%)
Differentiation time (hrs)
(D)
24 h
48 h
Myogenin-positive nuclei (%)
MyoD1 intensity (a.u.)
(E)
Percentage nuclei expressing Halo-MyoD1 (%)
1000 ng/mL
500 ng/mL
250 ng/mL
125 ng/mL
61 ng/mL
31 ng/mL
15 ng/mL
No Dox
(F)
Halo-MyoD1 fluorescence intensity (a.u.)
1000 ng/mL
500 ng/mL
250 ng/mL
125 ng/mL
61 ng/mL

Supplementary Figure 2

(A) Representative confocal microscopy images of Halo-MyoD1 expression and Myogenin (MyoG) immunohistochemistry. Images represent the C2C12$^{iMyoD1}$ cell line (Fig. 1A) treated ± dox for 24 or 48 h of differentiation. Yellow; JFX549 Halo-tag ligand-labeled Halo-MyoD1, Green; Myogenin (MyoG) immunohistochemistry, Blue; Hoechst labeled nuclei. Scale bar: 100 $\mu$m.

(B) Quantification of (A) representing the percentage of dox-treated C2C12$^{iMyoD1}$ nuclei expressing detectable Halo-MyoD1. Boxplots represent percentage of nuclei expressing Halo-MyoD1 above a background fluorescence intensity threshold. Plotted bars and whiskers represent 3 biological replicates. Analysis represents a minimum of 400,000 nuclei per time point and condition. Boxplots represent quartile ranges and medians, as detailed in Fig 1.

(C) Quantification of the percentage of nuclei expressing Myogenin (MyoG) based on their Halo-MyoD1 expression. Boxplots represent percentage of nuclei per well expressing MyoG above a background fluorescence intensity threshold. Bars and whiskers represent 3 biological replicates. Analysis represents a minimum of 400,000 nuclei per time point and condition.

(D) Percentage of Myogenin-positive nuclei as a function of Halo-MyoD1 expression level in C2C12$^{iMyoD1}$ cells at 24 h (blue line) and 48 h (orange line). Cells were binned by Halo-MyoD1 intensity, and points represent the mean Myogenin positivity within each bin. Whiskers indicate SEM over three biological replicates. Analysis includes a minimum of 80,000 nuclei per condition and time point.

(E) Quantification of the percentage of C2C12$^{iMyoD1}$ cells expressing Halo-MyoD1 after 24 h of differentiation and induction by a gradient of dox concentrations. Boxplots represent percentage of nuclei per well expressing Halo-MyoD1 above a background fluorescence intensity threshold. Bars and whiskers represent 3 biological replicates. Analysis represents a minimum of 200,000 nuclei per condition. No dox represents an untreated control.

(F) Violin plot quantification of the fluorescence intensity of Halo-MyoD1-expressing C2C12$^{iMyoD1}$ cells as a function of induction by a gradient of dox concentrations. Plotted bars represent median of nuclei pooled over 3 biological replicates. Analysis represents a minimum of 30,000 nuclei per condition.

### FIG-S3

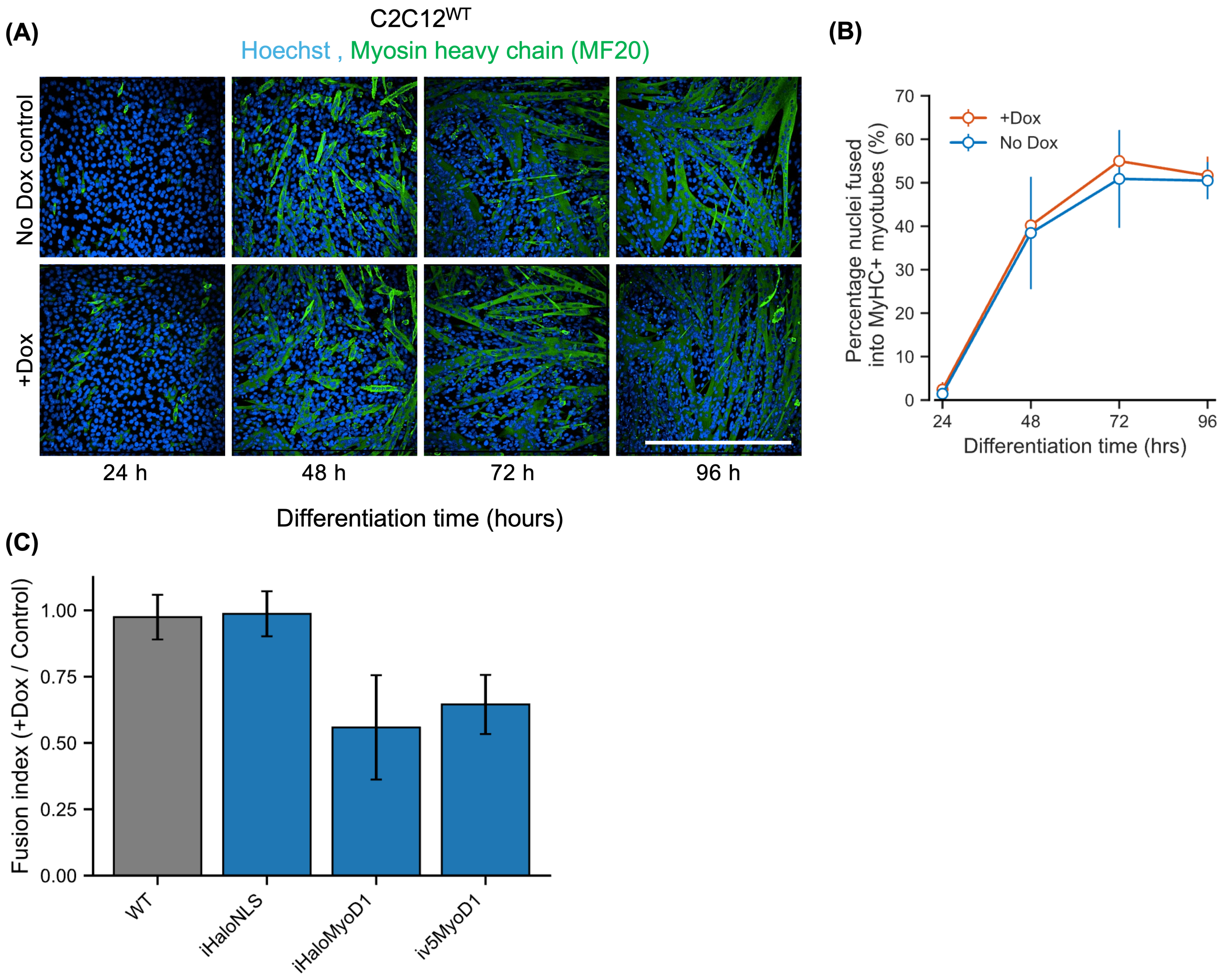

(A)
C2C12$^{WT}$
Hoechst , Myosin heavy chain (MF20)
No Dox control
+Dox
24 h
48 h
72 h
96 h
Differentiation time (hours)
(B)
70
60
50
40
30
20
10
0
+Dox
No Dox
Percentage nuclei fused into MyHC+ myotubes (%)
24
48
72
96
Differentiation time (hrs)
(C)
Fusion index (+Dox / Control)
1.00
0.75
0.50
0.25
0.00
WT
iHaloNLS
iHaloMyoD1
iv5MyoD1

Supplementary Figure 3

(A) Representative images of C2C12$^{WT}$ culture nuclei (Hoechst: blue) and myotubes (Myosin Heavy Chain immunofluorescence: green) over 96 h differentiation ± dox Scale bar: 500 $\mu$m.

(B) Quantification of C2C12$^{WT}$ ±dox myoblast fusion into MyHC+ myotubes. Plotted dots and bars represent means and standard deviation across 3 biological replicates.

(C) Quantification representing the ratio of fusion at 96 h differentiation. Fusion index represents the percentage of fused nuclei in +dox condition divided by no dox control. Analysis represents measurements of >450,000 nuclei over 3 biological replicates. Bars represent fusion index of C2C12$^{WT}$, C2C12$^{iHaloNLS}$, C2C12$^{iHaloMyoD1}$ and C2C12$^{iV5MyoD1}$. Error bars represent SEM of biological replicates.

### FIG-S4

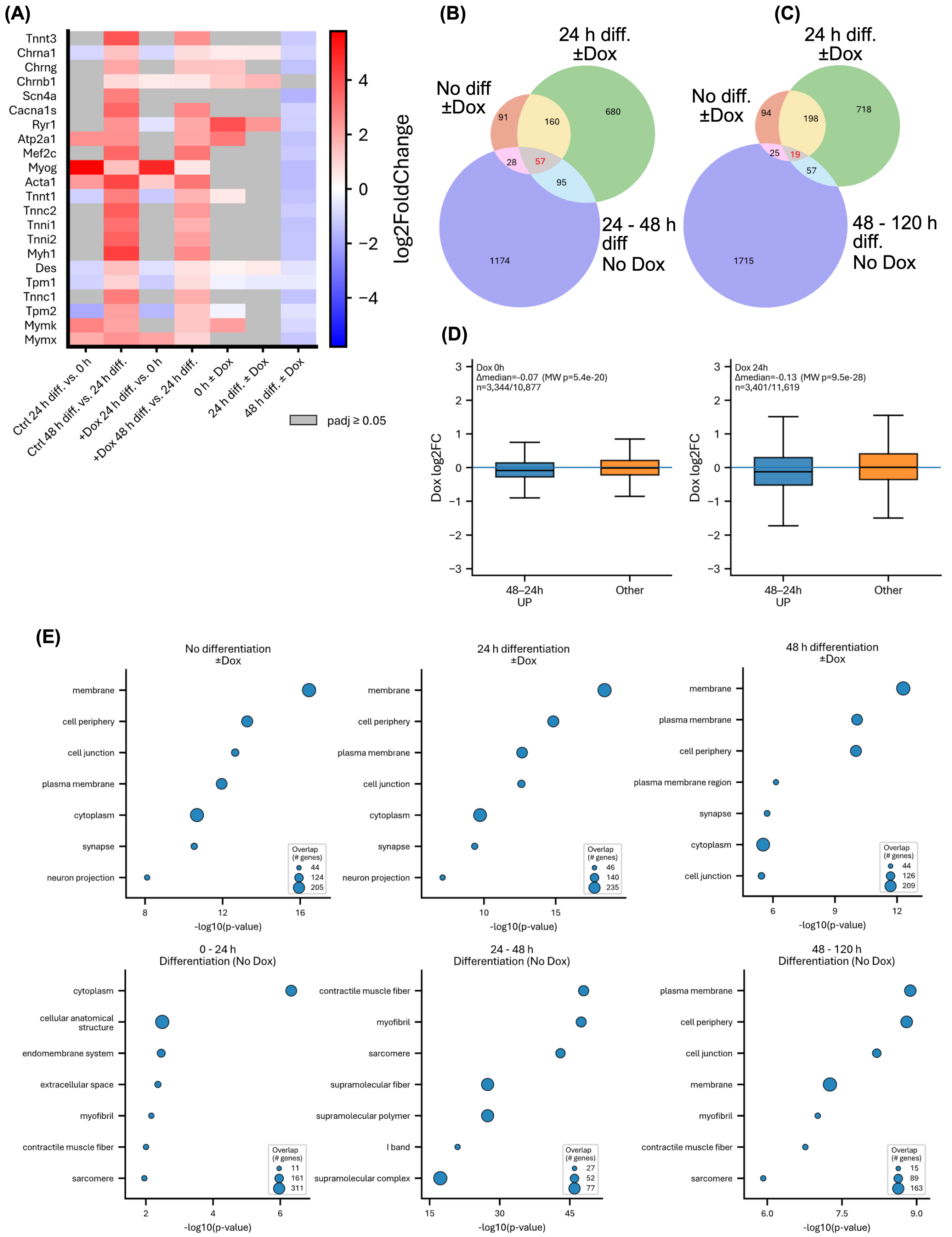

(A)
Tnnt3
Chrna1
Chrng
Chrnb1
Scn4a
Cacna1s
Ryr1
Atp2a1
Mef2c
Myog
Acta1
Tnnt1
Tnnc2
Tnni1
Tnni2
Myh1
Des
Tpm1
Tnnc1
Tpm2
Mymk
Mymx
log2FoldChange
Ctrl 24 h diff. vs. 0 h
Ctrl 48 h diff. vs. 24 h diff.
+Dox 24 h diff. vs. 0 h
+Dox 48 h diff. vs. 24 h diff.
0 h ± Dox
24 h diff. ± Dox
48 h diff. ± Dox
padj ≥ 0.05
(B)
24 h diff. ±Dox
No diff ±Dox
24 - 48 h diff No Dox
91
160
680
28
57
95
1174
(C)
24 h diff. ±Dox
No diff. ±Dox
48 - 120 h diff. No Dox
94
198
718
25
19
57
1715
(D)
Dox 0h
Δmedian=-0.07 (MW p=5.4e-20)
n=3,344/10,877
Dox 24h
Δmedian=-0.13 (MW p=9.5e-28)
n=3,401/11,619
Dox log2FC
48–24h UP
Other
(E)
No differentiation ±Dox
24 h differentiation ±Dox
48 h differentiation ±Dox
0 - 24 h Differentiation (No Dox)
24 - 48 h Differentiation (No Dox)
48 - 120 h Differentiation (No Dox)
membrane
cell periphery
cell junction
plasma membrane
cytoplasm
synapse
neuron projection
plasma membrane region
cellular anatomical structure
endomembrane system
extracellular space
myofibril
contractile muscle fiber
sarcomere
supramolecular fiber
supramolecular polymer
I band
supramolecular complex
Overlap (# genes)
-log10(p-value)

Supplementary Figure 4

(A) Heatmap of DESeq2 log2 fold-change values for a curated set of previously reported MyoD1-related genes involved in myogenic differentiation. Columns show pairwise comparisons across myogenic differentiation timepoints in control and dox-treated conditions, as well as matched-timepoint comparisons of dox-treated versus control cells. Cells are colored by DESeq2 log2 fold change for genes with significant differential expression between the indicated conditions; grey cells indicate non-significant changes. Red indicates higher expression in the numerator condition of each comparison, while blue indicates lower expression. Significance was defined as Benjamini-Hochberg adjusted $P < 0.05$. Columns correspond to comparisons: 0-24 h diff., 24-48 h diff., 0-24 h diff. +dox, 24-48 h diff. +dox and change due to dox treatment at 0 h diff. ±dox, 24 h diff. ±dox and 48 h diff. ±dox.

(B) 3-way Venn diagram of upregulated gene overlap (>2-fold, p.adj<0.05) between genes upregulated due to +dox treatment without differentiation (red circle.), +dox treatment with 24 h differentiation (green circle) and due to endogenous myoblast differentiation (24-48 h differentiation, no dox, blue circle). Number annotation represents the number of upregulated genes in each Venn category.

(C) 3-way Venn diagram as in (B) as compared to endogenous myoblast differentiation (48-120 h differentiation, no dox, blue circle).

(D) Boxplots of dox-induced log2 fold-changes (dox vs control) at 0 h (left) and 24 h (right) for genes significantly upregulated during endogenous differentiation from 48 h vs 24 h in control (48–24 h UP; padj < 0.05, log2FC > 0) compared with all other genes (“Other”). Center lines indicate medians; boxes indicate interquartile ranges; whiskers indicate 1.5×IQR, Δmedian and two-sided Mann–Whitney p-values are shown; n indicates the number of genes in each group.

(E) Cell compartment gene ontology analysis of up to top 400 significantly + dox upregulated (adj. p <0.05, 2-fold) genes in +dox C2C12$^{iMyoD1}$ (proliferating or simultaneous with 24 and 48 h differentiation) and top genes upregulated by endogenous differentiation (0-24, 24-48 and 48-120 h).

### FIG-S5

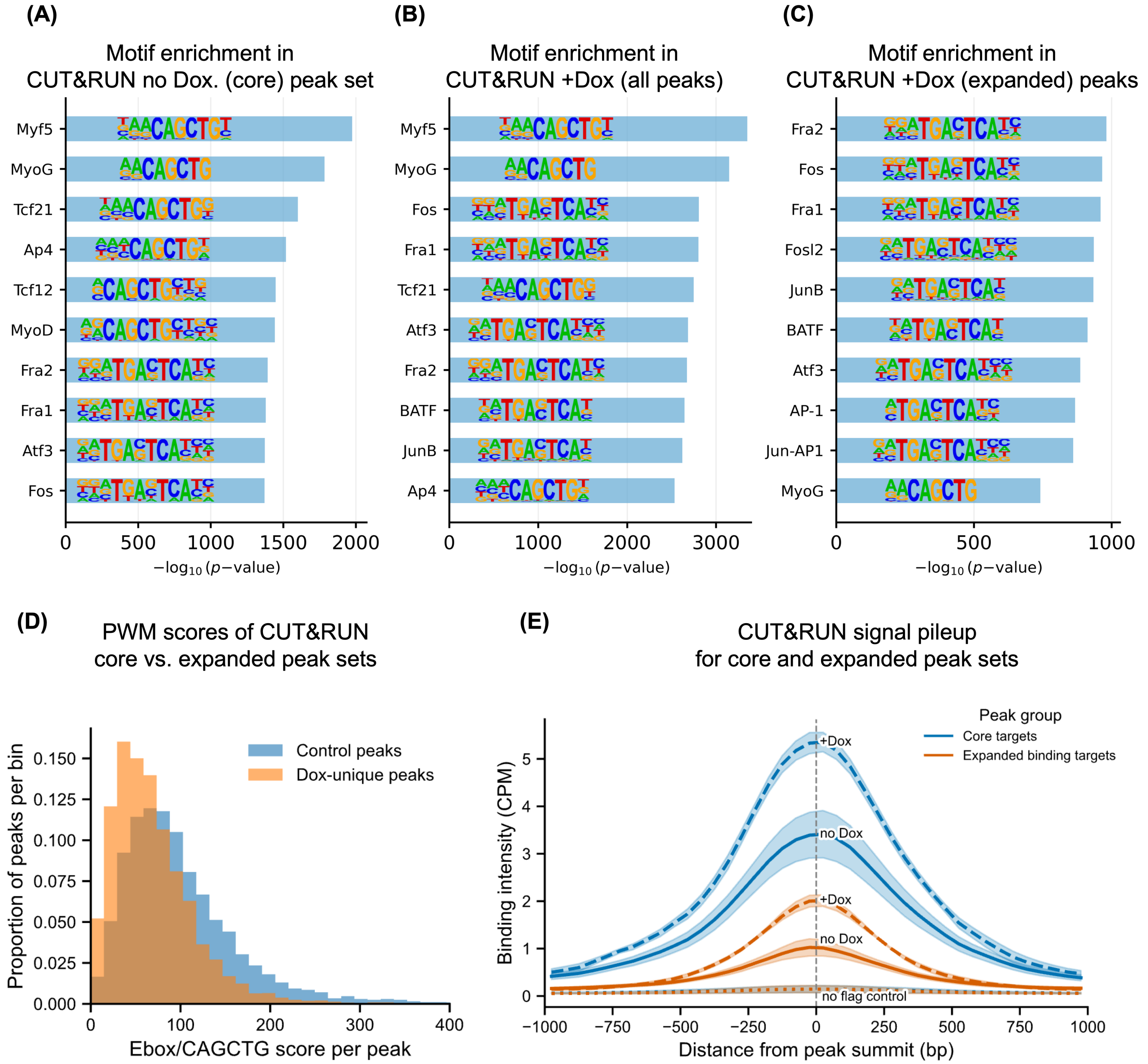
(A)
Motif enrichment in CUT&RUN no Dox. (core) peak set
Myf5
MyoG
Tcf21
Ap4
Tcf12
MyoD
Fra2
Fra1
Atf3
Fos
0
500
1000
1500
2000
−log10 (p−value)
(B)
Motif enrichment in CUT&RUN +Dox (all peaks)
Myf5
MyoG
Fos
Fra1
Tcf21
Atf3
Fra2
BATF
JunB
Ap4
0
1000
2000
3000
−log10 (p−value)
(C)
Motif enrichment in CUT&RUN +Dox (expanded) peaks
Fra2
Fos
Fra1
Fosl2
JunB
BATF
Atf3
AP-1
Jun-AP1
MyoG
0
500
1000
−log10 (p−value)
(D)
PWM scores of CUT&RUN core vs. expanded peak sets
Control peaks
Dox-unique peaks
Proportion of peaks per bin
0.000
0.025
0.050
0.075
0.100
0.125
0.150
0
100
200
300
400
Ebox/CAGCTG score per peak
(E)
CUT&RUN signal pileup for core and expanded peak sets
Peak group
Core targets
Expanded binding targets
+Dox
no Dox
+Dox
no Dox
no flag control
Binding intensity (CPM)
0
1
2
3
4
5
−1000
−750
−500
−250
0
250
500
750
1000
Distance from peak summit (bp)

Supplementary Figure 5

(A) Motif enrichment analysis of “core” CUT&RUN peak set vs mm10 background genome. Bars represent top 10 most significantly enriched motifs. Within-bar-lettering represents position-weight matrix of TF-associated motifs.

(B) Motif enrichment analysis of all dox-treated CUT&RUN peaks vs mm10 background genome, as in (A).

(C) Motif enrichment analysis of dox-unique CUT&RUN peak set vs mm10 background genome, as in (A).

(D) Ebox/CAGCTG motif PWM-scan scores for core (Control) and expanded (Dox-unique) CUT&RUN peak sets. Histogram represents distribution of peak motif scores within each condition. Orange distribution represents +dox-unique peaks, blue distribution represents no dox treatment (control) peaks.

(E) Per-condition pileup of CPM-normalized CUT&RUN signal under no-dox control (blue, core targets) or +dox-unique (orange, expanded binding targets) peak sets. +Dox-CUT&RUN signal indicated by dashed lines, no dox signal indicated by solid lines. No Flag control refers to CUT&RUN performed using anti-flag antibody in a parental C2C12 wild-type cell line not expressing any flag-tagged protein. Line shading represents ±1 standard deviation over three biological CUT&RUN replicates.

### FIG-S6

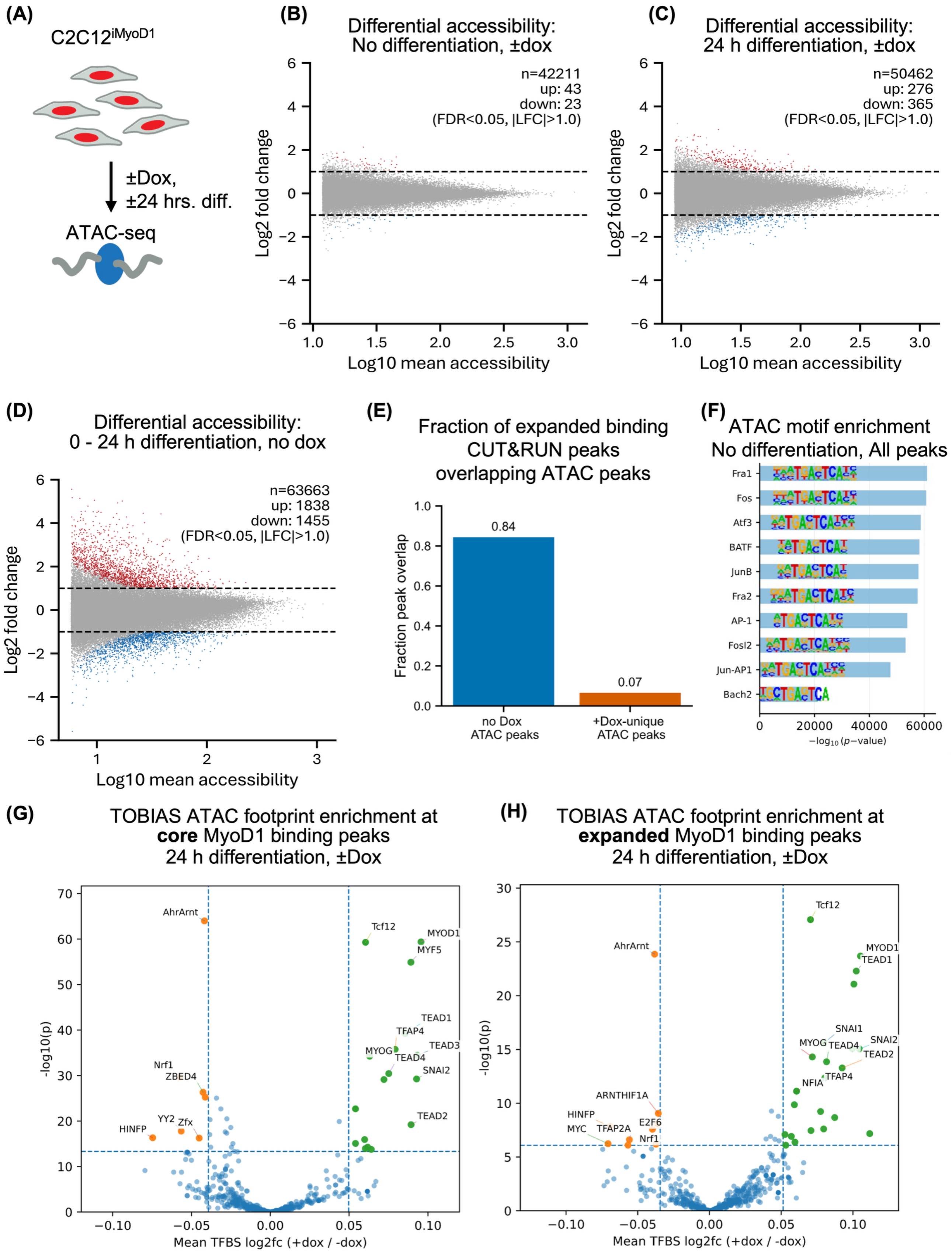

(A)
C2C12iMyoD1
±Dox,
±24 hrs. diff.
ATAC-seq
(B)
Differential accessibility:
No differentiation, ±dox
n=42211
up: 43
down: 23
(FDR<0.05, |LFC|>1.0)
Log2 fold change
Log10 mean accessibility
(C)
Differential accessibility:
24 h differentiation, ±dox
n=50462
up: 276
down: 365
(FDR<0.05, |LFC|>1.0)
Log2 fold change
Log10 mean accessibility
(D)
Differential accessibility:
0 - 24 h differentiation, no dox
n=63663
up: 1838
down: 1455
(FDR<0.05, |LFC|>1.0)
Log2 fold change
Log10 mean accessibility
(E)
Fraction of expanded binding
CUT&RUN peaks
overlapping ATAC peaks
Fraction peak overlap
0.84
0.07
no Dox
ATAC peaks
+Dox-unique
ATAC peaks
(F)
ATAC motif enrichment
No differentiation, All peaks
Fra1
Fos
Atf3
BATF
JunB
Fra2
AP-1
Fosl2
Jun-AP1
Bach2
−log10 (p−value)
(G)
TOBIAS ATAC footprint enrichment at
core MyoD1 binding peaks
24 h differentiation, ±Dox
AhrArnt
Tcf12
MYOD1
MYF5
TEAD1
TFAP4
TEAD3
MYOG
TEAD4
SNAI2
Nrf1
ZBED4
YY2
Zfx
HINFP
TEAD2
-log10(p)
Mean TFBS log2fc (+dox / -dox)
(H)
TOBIAS ATAC footprint enrichment at
expanded MyoD1 binding peaks
24 h differentiation, ±Dox
Tcf12
AhrArnt
MYOD1
TEAD1
SNAI1
MYOG
TEAD4
SNAI2
TEAD2
TFAP4
NFIA
ARNTHIF1A
HINFP
E2F6
MYC
TFAP2A
Nrf1
-log10(p)
Mean TFBS log2fc (+dox / -dox)

Supplementary Figure 6

(A) Schematic describing ATAC sequencing of C2C12$^{iMyoD1}$ ± Dox treatment and ± 24 h differentiation. Cells with red nuclei describe C2C12$^{iMyoD1}$, blue oval and wavy arms represent transposase and adapter oligos employed in ATAC-seq.

(B) Accessibility log-ratio to average (MA) plot of C2C12$^{iMyoD1}$ ATAC-seq. ± dox and no differentiation. Red dots indicate significantly (FDR<0.05), >2-fold more accessible peaks, blue dots indicate significantly (FDR<0.05), >2-fold less accessible peaks. Dashed lines represent a 2-fold change threshold. N_up and N_down represent the number of peaks up/down-regulated by 2-fold.

(C) (MA) plot of C2C12$^{iMyoD1}$ ATAC-seq. ± dox and 24 h differentiation, as in (B).

(D) (MA) plot of C2C12$^{iMyoD1}$ ATAC-seq. 0 – 24 h differentiation, as in (B).

(E) Fraction of overlap between +dox-unique MyoD1 CUT&RUN peaks and all ATAC peaks of either untreated or unique to +dox treated C2C12$^{iMyoD1}$. CUT&RUN and ATAC peaks represent SEACR (CUT&RUN) and MACS2 (ATAC) peaks reproducible in ≥2/3 biological replicates.

(F) Motif enrichment analysis of all ATAC peaks in untreated, proliferating C2C12$^{iMyoD1}$. Bars represent top 10 most significantly enriched motifs. Within-bar-lettering represents position-weight matrix of TF-associated motifs.

(G) Volcano plot of TOBIAS footprinting analysis across **core** binding-associated ATAC peaks in differentiating myoblasts ±dox (24 h diff.). Each point represents a TF footprint, x-axis shows mean TFBS log2 fold change in footprinting (±dox), y-axis shows -log10(*p*). Dashed lines indicate quantile-based cutoffs; labeled TFs are the strongest differential hits.

(H) Volcano plot of TOBIAS footprinting analysis across **expanded** binding-associated ATAC peaks, as in (G).

### FIG-S7

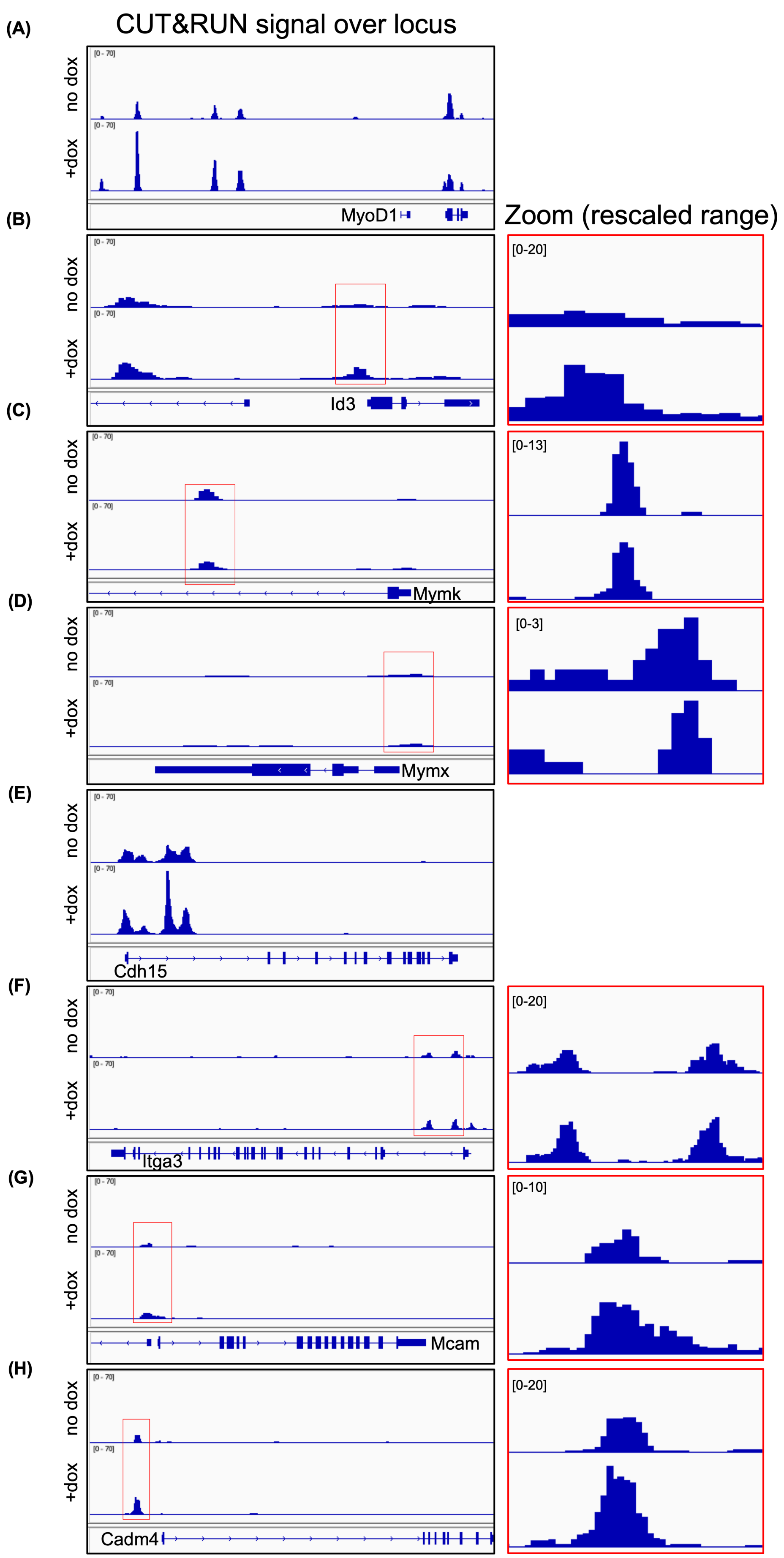

(A)
CUT&RUN signal over locus
no dox
+dox
[0 - 70]
MyoD1
(B)
Zoom (rescaled range)
Id3
[0-20]
(C)
Mymk
[0-13]
(D)
Mymx
[0-3]
(E)
Cdh15
(F)
Itga3
[0-20]
(G)
Mcam
[0-10]
(H)
Cadm4
[0-20]

Supplementary Figure 7

Example IGV tracks of Halo-MyoD1 CUT&RUN ±dox coverage at known targets and adhesion-related genes, including rescaled zoom for low occupancy targets.

(A) MyoD1 CUT&RUN coverage over the *MyoD1* locus (top lane is no dox, bottom is +dox)
(B) Coverage over *Id3* locus
(C) Coverage over *Mymk* locus
(D) Coverage over *Mymx* locus
(E) Coverage over *Cdh15* locus
(F) Coverage over *Itga3* locus
(G) Coverage over *Mcam* locus
(H) Coverage over *Cadm4* locus

#### FIG-S8

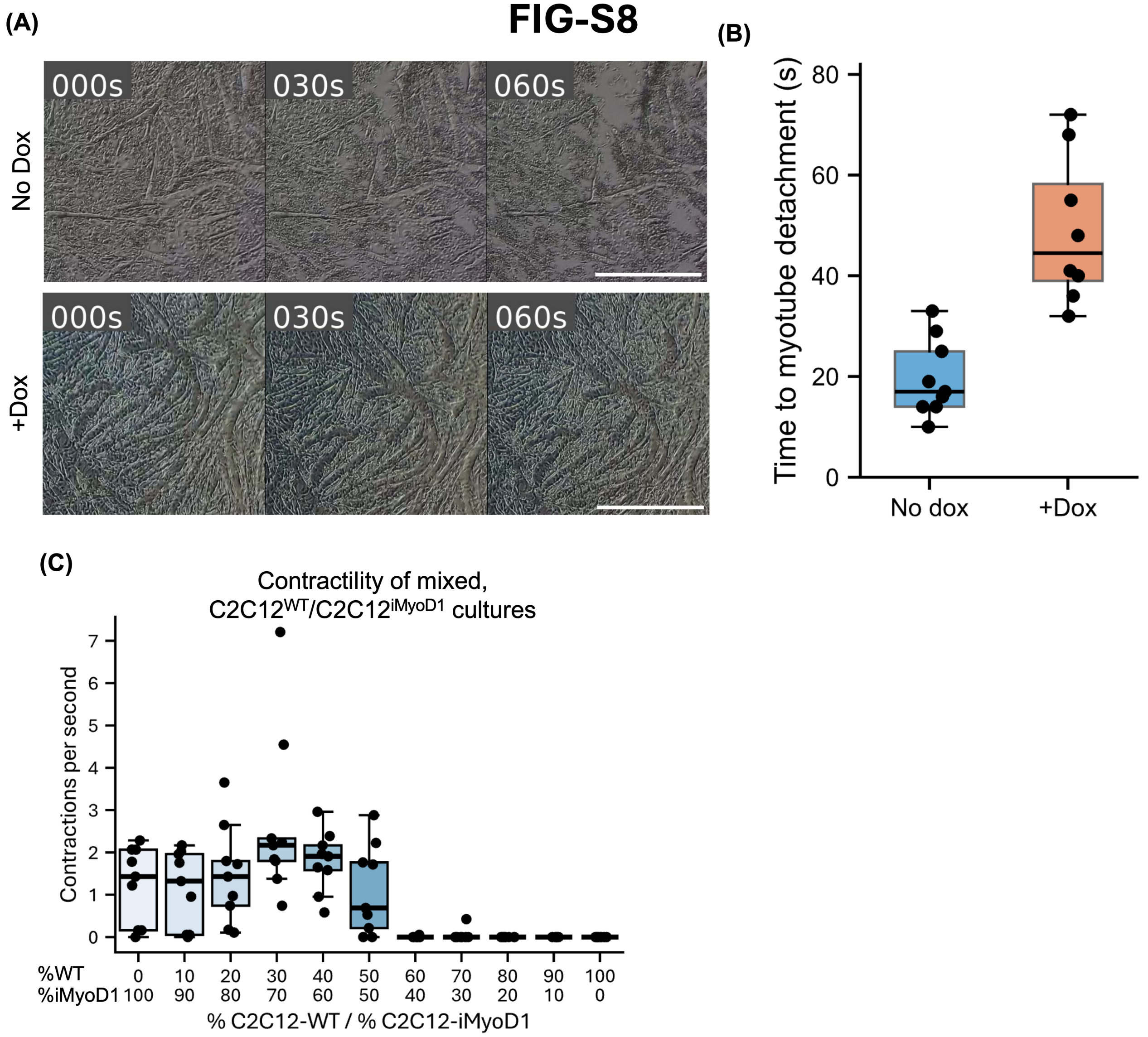
(A)
(B)
(C)
000s
030s
060s
No Dox
+Dox
Time to myotube detachment (s)
80
60
40
20
0
No dox
+Dox
Contractility of mixed,
C2C12WT/C2C12iMyoD1 cultures
Contractions per second
7
6
5
4
3
2
1
0
%WT
%iMyoD1
0 100
10 90
20 80
30 70
40 60
50 50
60 40
70 30
80 20
90 10
100 0
% C2C12-WT / % C2C12-iMyoD1

Supplementary Figure 8

(A) Snapshots of timelapse of trypsin treatment of C2C12$^{iMyoD1}$ cultures treated ±dox and differentiated for 120 h. Scale bar represents 200 $\mu$m.

(B) Quantification of (A), representing the time to initial myotube detachment after trypsin treatment addition. Dots represent ≥8 technical well replicates over 3 biological replicates.

(C) Optical flow contraction measurement quantification of mixed C2C12$^{iMyoD1}$/C2C12$^{WT}$ cultures treated +dox and differentiated 144 h. Dots represent 9 well replicates across 3 biological replicates.

FIG-S9

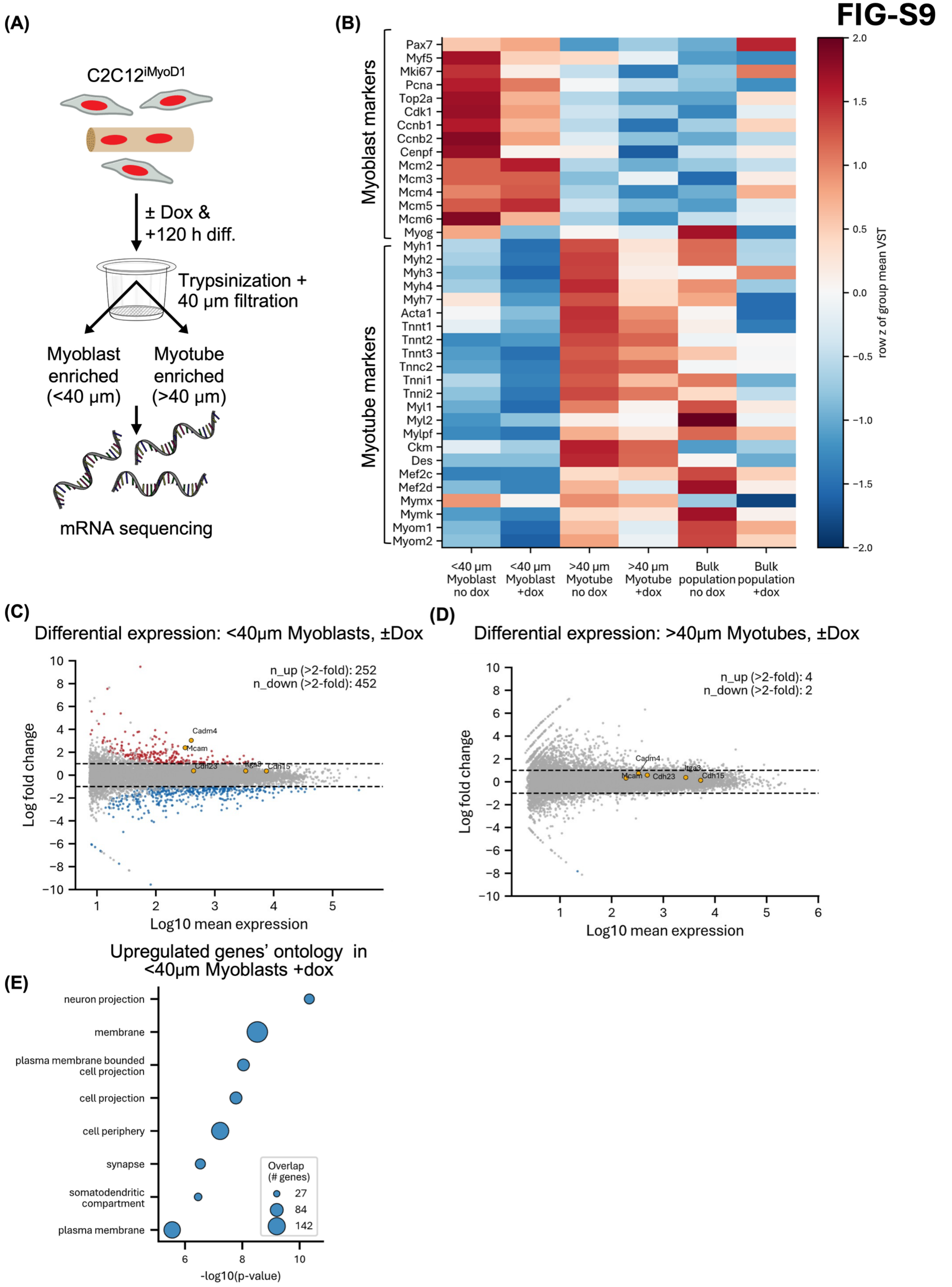
(A)
C2C12iMyoD1
± Dox &
+120 h diff.
Trypsinization +
40 μm filtration
Myoblast enriched (<40 μm)
Myotube enriched (>40 μm)
mRNA sequencing
(B)
Myoblast markers
Pax7
Myf5
Mki67
Pcna
Top2a
Cdk1
Ccnb1
Ccnb2
Cenpf
Mcm2
Mcm3
Mcm4
Mcm5
Mcm6
Myog
Myotube markers
Myh1
Myh2
Myh3
Myh4
Myh7
Acta1
Tnnt1
Tnnt2
Tnnt3
Tnnc2
Tnni1
Tnni2
Myl1
Myl2
Mylpf
Ckm
Des
Mef2c
Mef2d
Mymx
Mymk
Myom1
Myom2
<40 μm Myoblast no dox
<40 μm Myoblast +dox
>40 μm Myotube no dox
>40 μm Myotube +dox
Bulk population no dox
Bulk population +dox
row z of group mean VST
(C)
Differential expression: <40μm Myoblasts, ±Dox
n_up (>2-fold): 252
n_down (>2-fold): 452
Cadm4
Mcam
Cdh23
Itga3
Cdh15
Log fold change
Log10 mean expression
(D)
Differential expression: >40μm Myotubes, ±Dox
n_up (>2-fold): 4
n_down (>2-fold): 2
Cadm4
Mcam
Cdh23
Itga3
Cdh15
Log fold change
Log10 mean expression
(E)
Upregulated genes' ontology in <40μm Myoblasts +dox
neuron projection
membrane
plasma membrane bounded cell projection
cell projection
cell periphery
synapse
somatodendritic compartment
plasma membrane
Overlap (# genes)
27
84
142
-log10(p-value)

Supplementary Figure 9

(A) Schematic describing mRNA sequencing of C2C12$^{iMyoD1}$ following 120 h differentiation ± dox treatment and filtration. Red cells represent C2C12$^{iMyoD1}$, red nuclei in tan fibers represent myotube-fused nuclei. Myotubes were trypsinized and filtered through a 40 $\mu$m cell strainer, to generate a myoblast-enriched fraction (flowthrough) and myotube-enriched fraction (retentate).

(B) Heatmap of myogenic cell state marker gene expression in fractionated 120 h differentiated C2C12$^{iMyoD1}$ cultures. Cultures were separated into <40 $\mu$m myoblast-enriched and >40 $\mu$m myotube-enriched fractions, with or without dox treatment, and compared to a 120 h bulk population. Rows and annotations show curated proliferative/myoblast-like and myotube marker genes. For each gene, VST-normalized RNA-seq expression was averaged across biological replicates within each group and then row z-scored across the displayed 120 h groups. Red indicates higher expression relative to that gene's mean across groups, and blue indicates lower expression; values are capped at ±2. Columns are ordered as <40 $\mu$m myoblast no dox, <40 $\mu$m myoblast +dox, >40 $\mu$m myotube no dox, >40 $\mu$m myotube +dox, bulk population no dox, and bulk population +dox.

(C) Gene expression log-ratio to average (MA) plot of filter-isolated C2C12$^{iMyoD1}$ myoblast-fraction ± dox and 120 h differentiation. Red dots indicate significantly (p. adj<0.05) >2-fold upregulated genes. Blue dots indicate significantly (p. adj<0.05) >2-fold downregulated genes. Dashed lines represent a 2-fold change threshold. N_up and N_down represent the number of genes up/down-regulated as described above. Annotated genes represent adhesion-related candidates.

(D) Gene expression log-ratio to average (MA) plot of filter-isolated C2C12$^{iMyoD1}$ myotube-fraction ± dox and 120 h differentiation, as in (C).

(E) Cell compartment gene ontology analysis of significantly + dox upregulated (adj. p <0.05, 2-fold) genes in dox-treated C2C12$^{iMyoD1}$ myoblast fraction.

### FIG-S10

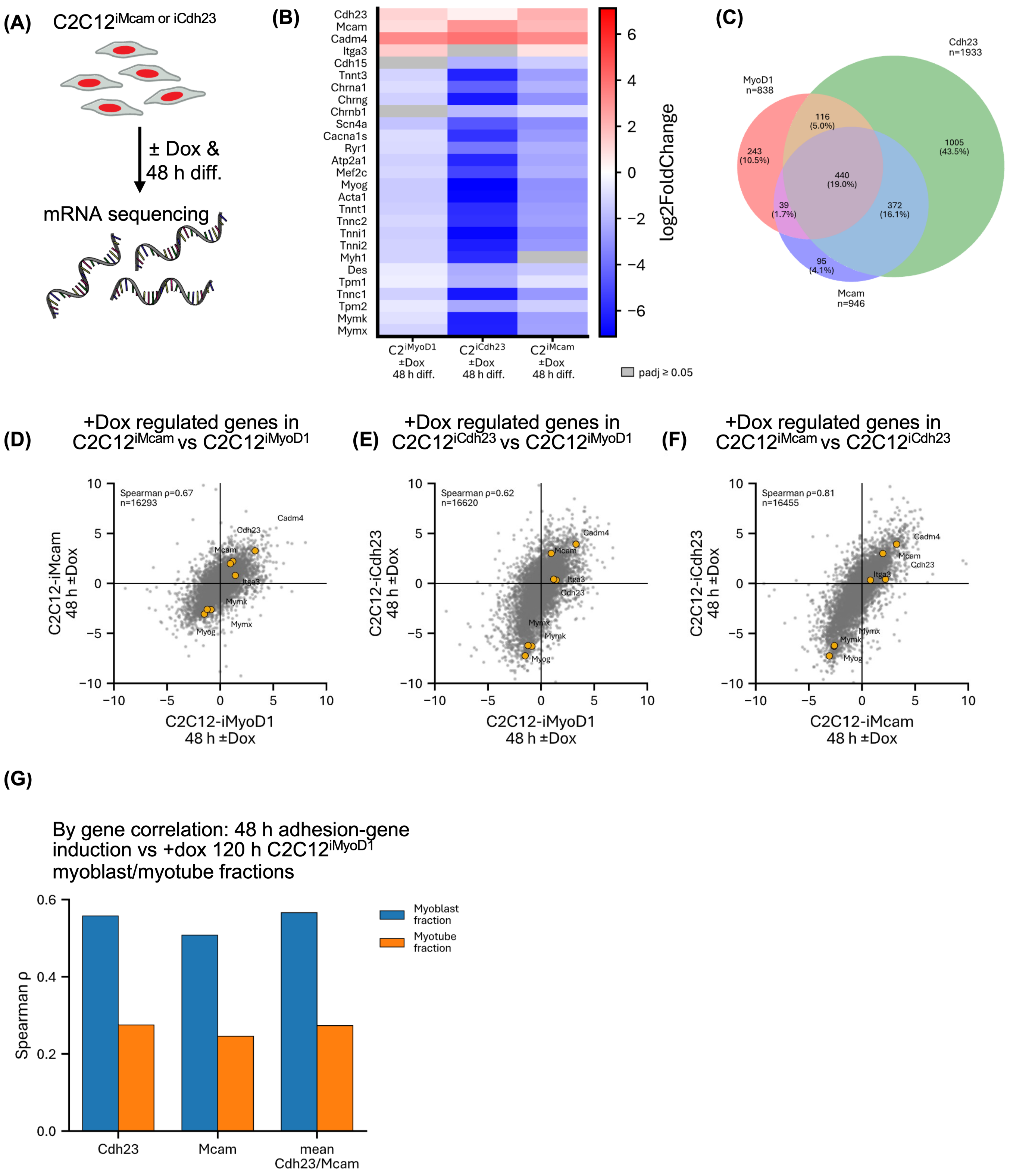
(A)
C2C12iMcam or iCdh23
± Dox &
48 h diff.
mRNA sequencing
(B)
Cdh23
Mcam
Cadm4
Itga3
Cdh15
Tnnt3
Chrna1
Chrng
Chrnb1
Scn4a
Cacna1s
Ryr1
Atp2a1
Mef2c
Myog
Acta1
Tnnt1
Tnnc2
Tnni1
Tnni2
Myh1
Des
Tpm1
Tnnc1
Tpm2
Mymk
Mymx
log2FoldChange
C2iMyoD1 ±Dox 48 h diff.
C2iCdh23 ±Dox 48 h diff.
C2iMcam ±Dox 48 h diff.
padj ≥ 0.05
(C)
Cdh23 n=1933
MyoD1 n=838
Mcam n=946
116 (5.0%)
243 (10.5%)
1005 (43.5%)
440 (19.0%)
39 (1.7%)
372 (16.1%)
95 (4.1%)
(D)
+Dox regulated genes in C2C12iMcam vs C2C12iMyoD1
Spearman ρ=0.67
n=16293
C2C12-iMcam 48 h ±Dox
C2C12-iMyoD1 48 h ±Dox
(E)
+Dox regulated genes in C2C12iCdh23 vs C2C12iMyoD1
Spearman ρ=0.62
n=16620
C2C12-iCdh23 48 h ±Dox
C2C12-iMyoD1 48 h ±Dox
(F)
+Dox regulated genes in C2C12iMcam vs C2C12iCdh23
Spearman ρ=0.81
n=16455
C2C12-iCdh23 48 h ±Dox
C2C12-iMcam 48 h ±Dox
(G)
By gene correlation: 48 h adhesion-gene induction vs +dox 120 h C2C12iMyoD1 myoblast/myotube fractions
Spearman ρ
Myoblast fraction
Myotube fraction
Cdh23
Mcam
mean Cdh23/Mcam

Supplementary Figure 10

(A) Schematic describing mRNA sequencing of C2C12$^{iMcam}$ or C2C12$^{iCdh23}$ following 48 h differentiation ± dox treatment. Red cells represent C2C12$^{iMcam/Cdh23}$.

(B) Heatmap of DESeq2 log2 fold-change values for a curated set of adhesion, myogenic differentiation, and fusion-related genes. Columns show within-cell-line comparisons of 48 h differentiated cultures treated with dox versus no dox in C2C12$^{iMyoD1}$, C2C12$^{iCdh23}$ and C2C12$^{iMcam}$. Cells are colored by DESeq2 log2 fold change for genes with significant differential expression between the indicated conditions; grey cells indicate non-significant changes. Red indicates higher expression with dox treatment, while blue indicates lower expression with dox treatment. Significance was defined as Benjamini-Hochberg adjusted $p < 0.05$.

(C) Three-way Venn diagram showing overlap among genes significantly upregulated following dox induction in C2C12$^{iMyoD1}$, C2C12$^{iCdh23}$, and C2C12$^{iMcam}$. Upregulated genes were defined as log2 fold-change > 1, *padj* < 0.05. Annotations indicate the number and percentage of upregulated genes.

(D) Gene-wise comparison of transcriptional responses to MyoD1 and Mcam induction. Scatterplot shows log2 fold-change values for +dox versus no dox treatment in C2C12$^{iMyoD1}$ cells and C2C12$^{iMcam}$ cells after 48 h differentiation. Each point represents one gene, labeled points indicate selected candidate adhesion, fusogen, and myogenic regulator genes. Spearman correlation coefficient and number of genes, annotated in top left.

(E) Gene-wise comparison of transcriptional responses to MyoD1 and Cdh23 induction, as in (D).

(F) Gene-wise comparison of transcriptional responses to Mcam and Cdh23 induction, as in (D).

(G) Spearman correlation between 48 h adhesion-gene induction and 120 h fractionated C2C12$^{iMyoD1}$ transcriptional responses. Bars show Spearman ρ values comparing gene-wise +dox versus no dox log2 fold-changes from 48 h C2C12$^{iCdh23}$, C2C12$^{iMcam}$, or the mean Cdh23/Mcam response to gene-wise +dox versus no dox log2 fold-changes from 120 h C2C12$^{iMyoD1}$ <40 $\mu$m myoblast-enriched or >40 $\mu$m myotube-enriched fractions.

Supplementary Video 1

(A) Real-time phase-contrast microscopy time-lapse of $C2C12^{iMyoD1}$ cells ± dox and 144 h of differentiation.

Supplementary Video 2

(A) Real-time phase-contrast microscopy time-lapse of $C2C12^{iCadm4}$, $C2C12^{iCdh15}$ $C2C12^{iCdh23}$, $C2C12^{iItga3}$, and $C2C12^{iMcam}$ cells ± dox and 216 h of differentiation.
